# Structural Basis of Herpesvirus Helicase-Primase Inhibition by Pritelivir and Amenamevir

**DOI:** 10.1101/2025.05.15.654119

**Authors:** Andrey G. Baranovskiy, Qixiang He, Yoshiaki Suwa, Lucia M. Morstadt, Nigar D. Babayeva, Ci Ji Lim, Tahir H. Tahirov

## Abstract

Widespread herpesvirus infections are associated with various diseases. DNA replication of human herpes simplex virus type 1 (HSV-1) requires a helicase-primase (HP) complex of three core proteins: UL5, UL52, and UL8. This complex unwinds viral DNA and synthesizes primers for DNA replication, making it an attractive antiviral target. Although HP inhibitors pritelivir and amenamevir were identified through screening, their binding mechanisms remain unclear. Here, we report cryo-EM structures of HSV-1 HP bound to a forked DNA template alone, and in complex with pritelivir or amenamevir. The structures reveal a bilobed architecture highlighting helicase-primase coordinated action at the replication fork and providing a structural basis for HP inhibition by illustrating precisely how pritelivir and amenamevir block helicase activity. Data lays a solid foundation for the development of improved antiviral therapies.

## Introduction

Herpesviruses are ubiquitous double-stranded DNA (dsDNA) viruses which can infect a wide range of hosts, including humans. Among them, herpes simplex virus 1 (HSV-1), one of eight known human herpesviruses, can establish lifelong infections in human cells through alternating lytic and latent cycles of infection. HSV-1 is associated with a variety of diseases including keratitis and encepohalitis.(*1, 2*) The linear, double-stranded HSV-1 genome is about 152 kilobases (kb) long and its genome DNA replication in infected human cells is initiated by a viral replication machinery which consists of six essential proteins.(*3*) These proteins are a single-stranded (ss) DNA-binding protein ICP8, a two-subunit DNA polymerase complex comprised of polymerase UL30 and processivity UL42 subunits, and a heterotrimeric helicase-primase (HP) complex comprised of helicase UL5, primase UL52, and auxiliary UL8 subunits.(*4, 5*)

After the origin binding protein UL9 binds to the replication origins, the HP complex (UL5-UL52-UL8) is recruited to the replication fork to unwind the viral duplex DNA (helicase activity) and build short RNA primers (primase activity) for DNA polymerase (UL30-UL42) to start leading and lagging strand DNA synthesis.(*4, 5*)

Biochemical studies showed that purified HSV-1 HP complex assembles as a 1:1:1 heterotrimer with a total molecular weight of 293 Kilodaltons (kDa).(*5*) The helicase UL5 belongs to the helicase superfamily 1 (SF1) and contains seven conserved sequence motifs critical for NTP-powered single-stranded DNA translocation activities (**Fig 1A**).(*6, 7*) The primase UL52 harbors three conserved motifs characteristic of archaeal-eukaryotic primase (AEP) catalytic domain, along with a putative zinc-binding domain that may facilitate DNA template binding (**Fig 1A**).(*8, 9*)

**Figure 1.**
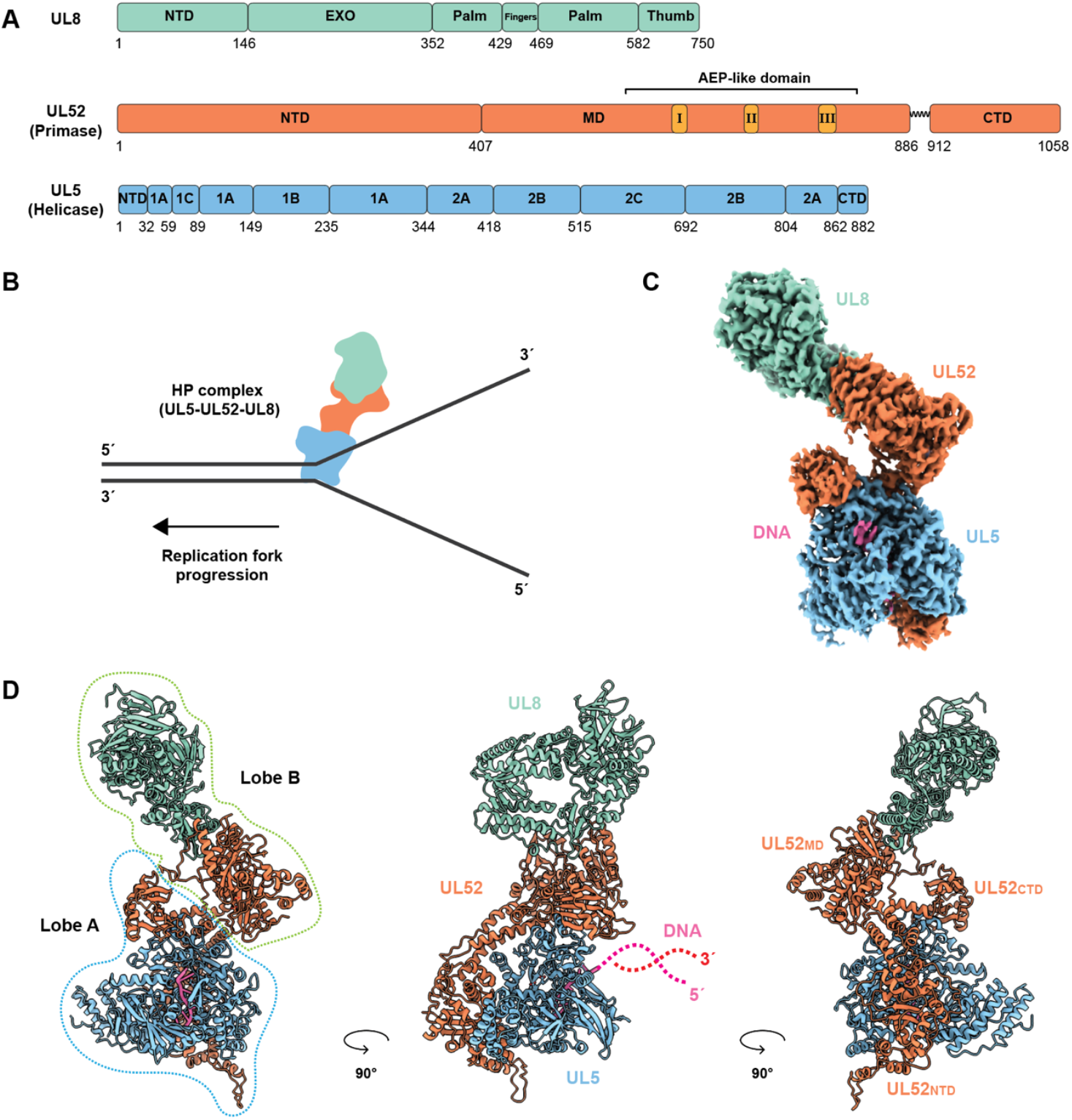
Overall structure of the HSV-1 HP complex bound to a forked DNA. (**A**) Domain organization of UL8, UL52 and UL5. NTD, N-terminal domain; EXO, exonuclease domain; MD, middle domain; CTD, C-terminal domain; AEP-like domain, archael-eukaryotic primase (AEP)-like domain. The conserved AEP motifs I-III in UL52 are shown in different color. (**B**) Schematic drawing of an HSV-1 HP complex at forked DNA. (**C**) Cryo-EM density map of the HP-complex bound a forked DNA substrate, colored by subunit. (**D**) Ribbon diagrams of the HP complex structure, also colored by subunit. Three views are shown for clarity. The dashed lines indicate the position of the duplex region of the forked DNA substrate, which was poorly resolved in density map.

Although UL5 and UL52 are respectively classified as the helicase and primase subunit of HP based on motifs homology, neither protein alone exhibits helicase or primase activity.(*10*) Their heterodimer assembly is needed to activate their helicase and primase activity, suggesting a tight interdependency between the two enzymes to mediate their respective functions.(*11-13*) UL8, which structurally resembles a B-family polymerase, is vital for optimal HP complex activity (**Fig 1A**).(*14, 15*) As UL8 possesses no known intrinsic catalytic activity, it is likely that its HP stimulatory function stems from its participation as a key architectural component.

The HP complex is an attractive target for antiviral drug development, as its DNA unwinding and primer synthesis functions are important precursory steps to processive DNA polymerization by the HSV DNA polymerase. Two HP inhibitors (HPIs), pritelivir(*16*) and amenamevir(*17*) (**Fig. S1**), were discovered through large-scale compound screening. Despite their efficacy, the mechanisms by which they inhibit HP remain unclear due to the lack of structural insights into their binding modes and how they disrupt enzymatic activity.

Here, we present cryo-electron microscopy (cryo-EM) structures of the HSV-1 helicase-primase (HP) complex bound to a forked DNA template and either pritelivir or amenamevir. These structures reveal the overall architecture of the HP complex when engaged with DNA. More importantly, the inhibitor-bound structures identify the binding sites of pritelivir and amenamevir and provide insights into how these compounds stall HP enzymatic activity.

### Overall architecture of HSV-1 HP complex bound to a forked DNA

To investigate the structural basis for the HP complex assembly at a replication fork, we expressed and purified the HP complex from insect cells for structural studies (**Fig. S2**). Previous DNA-binding assays(*18, 19*) and our study here (**Fig. S3**) indicate a preferential binding of HP complex to a forked DNA template with a 3’-flap. To gain structural insights into a template-bound HP catalytic state, we used cryo-EM single-particle analysis to capture the structure of HSV-1 HP complex bound to the forked DNA with a 3’-flap (**Figs S4, S5 and S6**). The resolved cryo-EM density maps confident docking of AlphaFold models for UL5, UL52, and UL8 (**Figs 1D, S4, S5 and S6**). Due to flexibility in the parental duplex region of the forked DNA, only seven nucleotides (nt) of the 3’-flap ssDNA—bound by UL5—were resolved and modeled. (**Figs 1D, S4, S5 and S6**).

The HP complex structure revealed a bilobed architecture with a 1:1:1 stoichiometry of UL52, UL5, and UL8 subunits (**Fig. 1C, 1D**). In this structure, the N-terminal domain of the UL52 (UL52_NTD_) and C-terminal domain of the UL52 subunit (UL52_CTD_) partially wrap around UL5 to form a compact, globular lobe (Lobe A) (**Fig. 1C**). The middle domain of UL52 (UL52_MD_) interacts with UL8 to form an elongated and flattened shape (Lobe B) (**Fig 1D**). These inter-subunit interactions are consistent with previous co-immunoprecipitation (co-IP) experiments.(*20*)

### UL52 DNA primase

UL52 primase subunit is extended along the molecule and acting as a scaffold for the HP complex assembly. UL52 has an elongated shape with three distinct domains: N-terminal (NTD), middle (MD), and C-terminal (CTD) domains (**Figs 2 and S7**). The NTD domain (residues 1-408), which lacks catalytic activity, contains a disordered 14-residue N-terminal tail. The remainder acquires a crescent-like shape with a novel fold without any similarities, as confirmed by DALI search.(*21*) It has a four-strand β-sheet close to its center, which contains two parallel β-strands in the middle and the outer β-strands antiparallel to the middle strands. It is surrounded by α-helices and two prominently protruding helical bundles (**Fig. 2A**).

**Figure 2.**
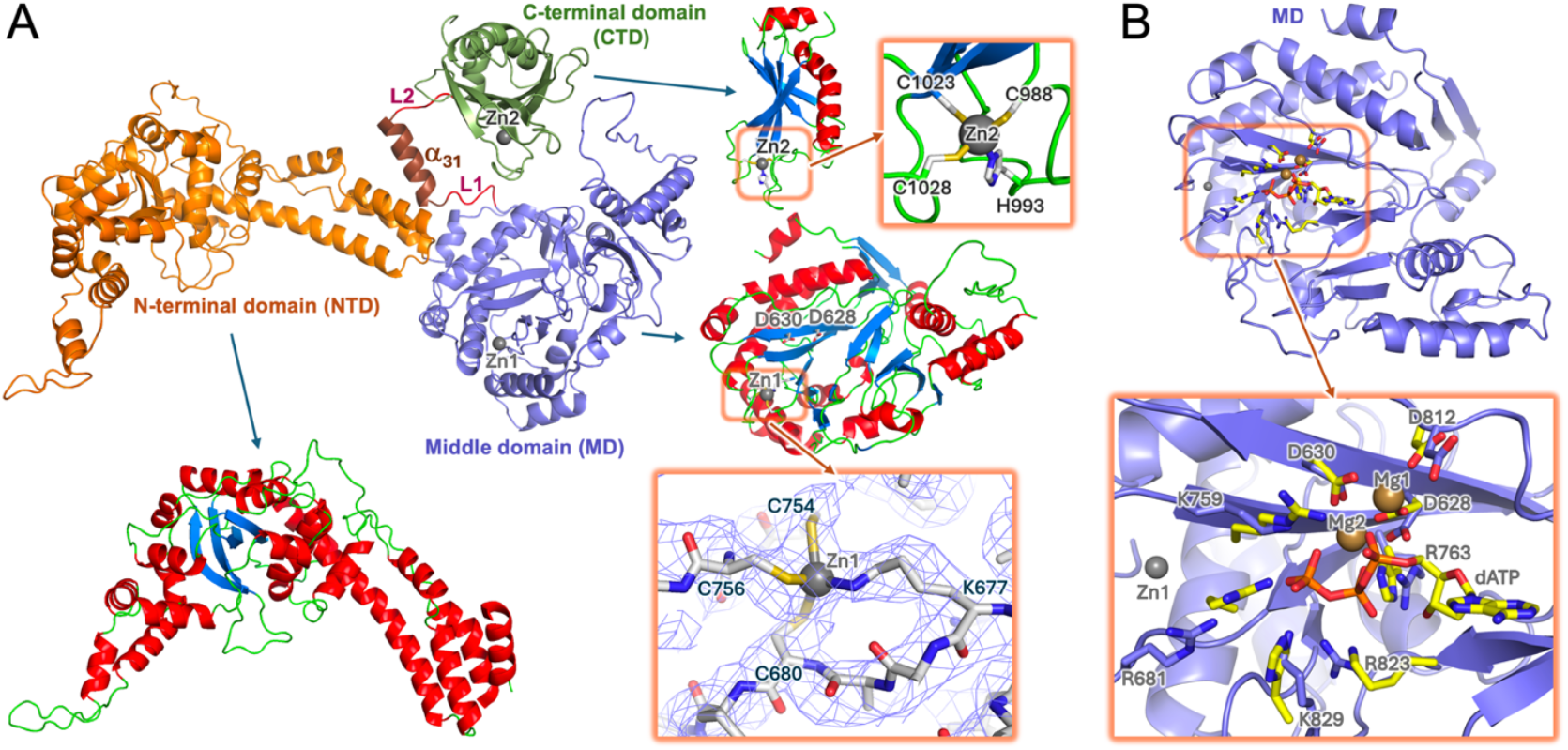
Structure of the HSV1 HP UL52 subunit. (**A**) Cartoon representation of UL52 colored by domains: NTD in orange, MD in slate, CTD in split pea green, linkers L1 and L2 in firebrick, and the helix α_31_ in brown. Each domain is displayed separately to highlight positions of α- and 3_10_-helices (red), *β*-strands (cyan), and coils (green). Zn^2+^ ions in CTD and MD are drawn as grey balls. Close-up views of each Zn^2+^ coordination are also shown. For Zn^2+^ in MD, the cryo-EM map is also displayed. (**B**) Primase active site in MD. The side chains of the amino acid in the active site are drawn as sticks and labeled in a close-up view. Crystal structure of the human primase Pri1 subunit in complex with two Mn^2+^ ions and the incoming nucleotide dATP (PDB code 6r5d)(*26*) has been superimposed with the MD of UL52 (rmsd 0.6 Å). The dATP, two Mn^2+^ ions, and the amino acid residues in the active site of the docked Pri1 are shown as yellow sticks.

Unlike the long flexible linker between the catalytic and C-terminal domains seen in the eukaryotic DNA primases, the connection between the corresponding domains in HSV1 primase contains a linker(L1)-α-helix(α_31_)-linker(L2) structure (**Fig. 2A**). The α_31_ helix (residues 893-907) in this linker is well-stabilized by interactions with an NTD. As a result, the potential movements of the CTD are limited by a four-residue linker, L2. Meanwhile, the relative position of the N-terminal and middle domains of UL52, and consequently the relative position of the HP Lobes A and B, is stabilized mainly by the inverse γ-turn centered at L408 and a linker L1 (**Fig. 2A**).

The MD domain (residue 409-887) corresponds to the catalytic domain found in the family of archaea-eukaryotic primases (AEP), which are characterized by the presence of three signature motifs, motifs I-III (**Fig. 1A and S7**).(*22*) The positions of the conserved residues from the corresponding motifs in HSV-1 HP are well aligned with that of human primase, PrimPol, and archaeal pRN1.(*23-25*) Superimposition of the high-resolution crystal structure of the human primase Pri1 subunit in complex with two Mn^2+^ ions and the incoming nucleotide dATP (PDB code 6r5d)(*26*) suggested possible UL52 binding pockets for ATP and two Mn^2+^ ions (**Fig. 2B**). Specifically, UL52 residues D628 and D630 in motif I, which are critical for the catalytic activity of primase(*8*), and D812 in the motif III most likely coordinate the catalytic metal ions, while R763 from motif III is poised to interact with the incoming nucleotide (**Fig. 2B**). R681, K759, R823, and K829 are also likely involved in binding to an incoming nucleotide.

Unlike the centrally located active site, the rest of the AEP catalytic domains are not conserved. The UL5 MD differs from the catalytic domain of human PrimPol and archaeal pRN1 which have relatively compact structures (surface area of 18788 Å^2^ vs 13747 Å^2^ and 10407 Å^2^, respectively) (**Fig. S8**). The human Pri1 is narrower but elongated as compared to the MD of UL5 (surface area of 18149 Å^2^ vs 18788 Å^2^), in part due to the presence of a helical bundle in the human Pri1 that is protruding in the direction where the N-terminal domain of UL52 is located (**Fig. S8**). The subunits interacting with the UL52 MD and human Pri1 (UL8 and Pri2, respectively) are located on the same side of the molecule, but their mode of interaction and the structures of the involved areas differ dramatically (**Fig. S8**). Similar to Pri1 and pRN1, the UL52 MD also contains a Zn^2+^ (**Fig. 2A**). However, unlike the Zn^2+^ in Pri1 or pRN1 (which are coordinated by four cysteines, or three cysteines and histidine, respectively), the Zn^2+^ in UL52 is coordinated by three cysteines (C680, C754, and C756) and a lysine (K677) (**Fig. 2A**). Such tetrahedral Zn^2+^ coordination with the involvement of lysine is rare. Nevertheless, a search conducted with MESPEUS, a database of the geometry of metal sites in proteins(*27*), confirmed the presence of similar zinc ion coordination in proteins.

The CTD domain (residue 912-1058) resembles the predicted structure of the Zn^2+^-containing domain of human PrimPol(*28*). The UL52 CTD is comprised of an antiparallel 5-strand β-sheet with an elongated banana-shape α-helix laying across the β-sheet (**Fig. 2A**). It also contains a Zn^2+^-binding site and a second shorter α-helix on the opposite ends of the domain. This CTD also contains a disordered C-terminal tail (1,046-1,058). The Zn^2+^ is coordinated by H993, C988, C1023, C1028 (Fig. 2A). Similar to the well-characterized CTDs of human primase Pri2 subunit(*29*) and the human PrimPol(*28*), the CTD of UL52 is a key contributor to binding of the DNA template and initiation of primer synthesis. Indeed, previous studies showed C1023A/C1028A substitutions disrupt the binding of Zn^2+^, which impacted the stability of the CTD fold and led to a reduction in the DNA-binding activities of the HP complex. This eliminated the primase activity, and, consequently, viral replication.(*13, 30*)

### UL5 helicase

UL5 is a Helicase Superfamily 1 (SF1) helicase possessing seven conserved motifs (**Figs. 3A and S9**).(*31*) It uses ATP or GTP hydrolysis to drive template translocation in a direction from 3’ to 5’, resulting in DNA duplex unwinding.(*19*) Cryo-EM studies of HP show a globular structure of UL5 containing all core elements of a four-domain (domains 1A, 1B, 2A, and 2B) SF1 helicase structure (**Fig. 3B**). However, UL5 exhibits notable topological differences and is significantly bulkier than most SF1 helicases. For example, among the structurally characterized members of the SF1 family, the closest in shape to UL5 is *Sc*Pif1.(*32*) Nevertheless, UL5 is significantly larger (surface area of 41,480 Å^2^ vs 25,249 Å^2^) and possesses several additional structural elements, mainly at the outer layer of the protein (**Fig. 3B**).

**Figure 3.**
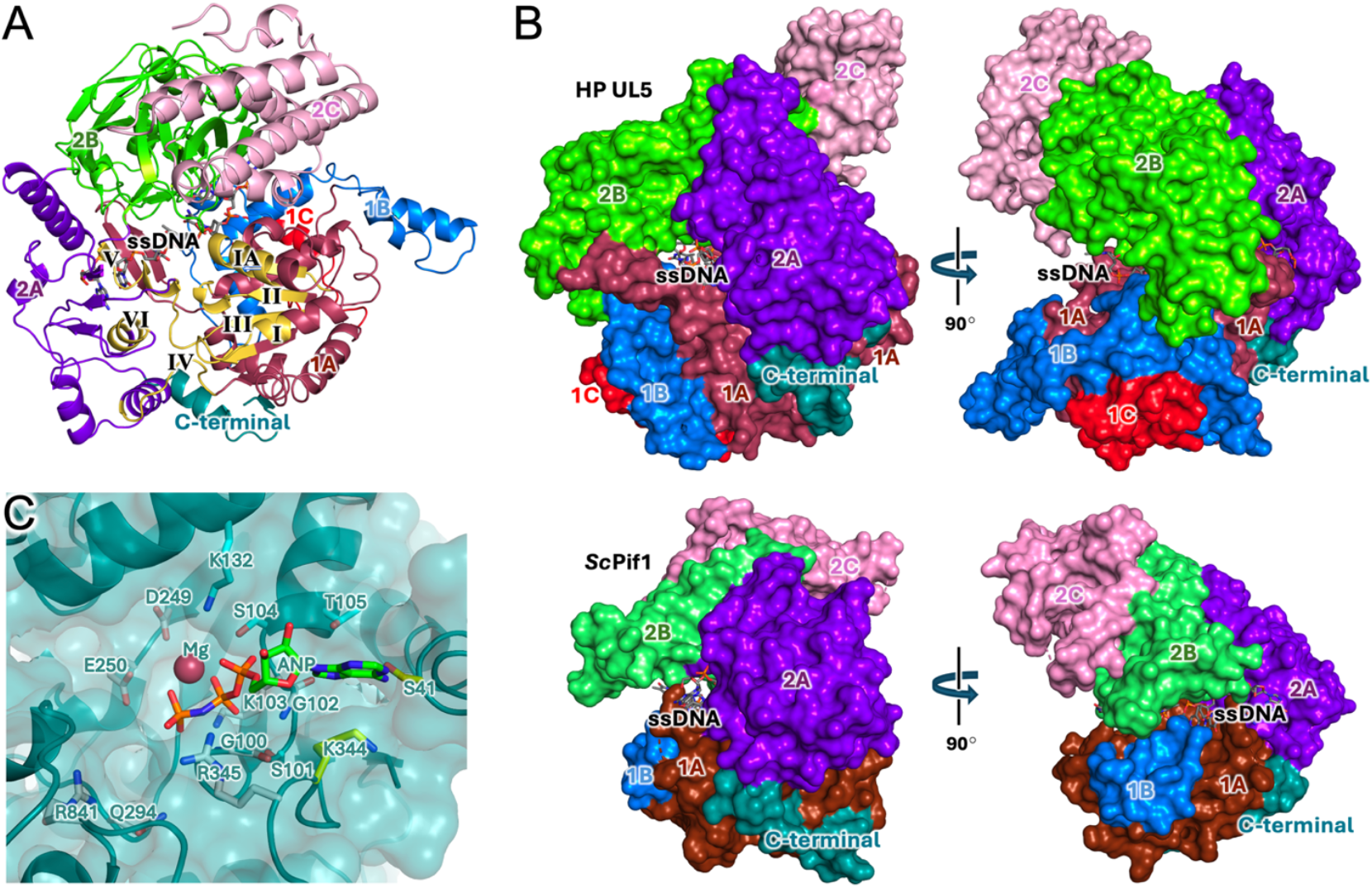
Structure of HSV1 HP UL5 subunit. (**A**) Cartoon representation of UL5 colored by domains: 1A in brown, 1B in cyan, 1C in red, 2A in purple, 2B in pale green, 2C in pink, and C-terminal in forest green. The helicase conserved motifs I-VI are highlighted in yellow. (**B**) Surface representations of HSV1 UL5 and *Sc*Pif1. The ssDNAs in both structures are represented as sticks and their domains are colored as in panel A. (**C**) Active site of UL5. The amino acid residues (G100, G102, K103, D249, E250, Q294, R345, R841) that are conserved in all SF1 helicases are drawn as white sticks. The residues that are conserved only among herpesviruses (S41 and K344) are drawn as yellow sticks. The non-conserved residues (S101, S104, T105, K132) are drawn as cyan sticks. To model the position of ADPNP and Mg^2+^ in UL5, the conserved motifs in domains 1A of UL5 and RecD2/ssDNA/ADPNP (PDB code 3gpl)(*33*) were aligned and then the coordinates of ADPNP and Mg^2+^ were merged with the coordinates of UL5.

One of the most notable differences is in domain 1B, which has an extended 5-helical structure that partially covers the surface of domain 1A, as compared to a single helix in ScPif1 (**Fig. 3B**). Moreover, domain 1B is additionally stabilized by a complementary two-helical insertion in domain 1A that lays on top of 1B and is further referred as domain 1C (**Fig. 3B**). Another domain showing the largest discrepancies is domain 2B. In UL5, domain 2B can be subdivided into two subdomains, one comprising a prominent *β*-sheet with five antiparallel strands and two α-helices, and another containing the SH3 domain with an insertion that includes two α-helices. In comparison, the domain 2B in *Sc*Pif1 contains a smaller three-strand *β*-sheet with one α-helix and an SH3 domain lacking the helical insertion (**Figs. 3B and S9**). Like *Sc*Pif1, UL5 also contains the helical domain 2C. However, the arrangement of the α-helices differs between the two proteins (**Fig. 3B**). Domain 2C in UL5 also contains a large 53-residue insertion that was poorly resolved in the cryo-EM maps (**Figs. S6, S10, and S11**) thus it is likely flexible or adopts multiple conformations.

The conserved helicase motifs are located at the interface of domains 1A and 2A: motifs I, IA, II, and III in domain 1A, motifs V and VI in domain 2A, and motif IV in the linker connecting domains 1A and 2A (**Fig. 3A**). Our cryo-EM structures of HP complex do not contain a bound ATP analog. Therefore, to locate and characterize the ATP/GTP binding site, we used the reported crystal structures of SF1 helicases as a reference. For example, the conserved motifs in domains 1A of UL5 and RecD2/ssDNA/ADPNP (3gpl,(*33*)) are aligned with a R.M.S.D. of 0.97 Å^2^ (for 47 α-carbons).

The residues in UL5 located in ATP/GTP and the Mg^2+^-binding area can be assigned into three groups (**Fig. 3C**). First are the amino acid residues that are conserved across all SF1 helicases (G100, G102, K103, D249, E250, Q294, R345, R841). Mutations in some of these residues have been shown to impair either ATP hydrolysis, DNA unwinding, or coupling between hydrolysis and unwinding.(*34*) Second are the residues that are conserved specifically among herpesviruses helicases (S41 and K344). K344 provides stacking for the NTP base, while S41 accommodates and directly interacts with either the ATP or GTP base, but not with CTP or TTP, and therefore, determines UL5 specificity for NTPs.(*19*) Excluding herpesvirus helicases, the majority of SF1 helicases are ATP-dependent and contain asparagine instead of serine (S41) and tyrosine or phenylalanine instead of lysine (K344). Their tyrosine or phenylalanine side chain stacks parallel to the ATP base and positions it for making two perfect H-bonds with asparagine, leading to high specificity toward ATP. Finally, the third group are the non-conserved residues (S101, S104, T105, K132). All these residues are expected to form H-bonds with the triphosphate moiety of ATP or GTP. Additionally, T105 provides a complementary surface for ATP base stacking on the side opposite from K344 (**Fig. 3C**).

Our DNA-binding experiments confirmed that the HP complex preferentially binds to a forked DNA at the 3’-overhang (**Fig. S3**).(*18, 19*) Indeed, our cryo-EM single particle analysis resolves seven nucleotides of the 3’-overhang (**Fig. S12A**). A DNA-binding channel runs close to the middle of UL5 and is surrounded by domains 1A, 1B, 2A, and 2B (**Fig. 3A**). Within this channel, thirteen potential H-bonds are observed between UL5 and the phosphate groups of ssDNA, and four potential H-bonds are formed involving ssDNA bases (**Fig. S12B**). Although the density for the duplex region is at low-resolution, it spans approximately seven base pairs, allowing us to infer the positioning of the duplex portion of forked DNA. The forked end of the duplex is leaning onto the surface of UL5 domain 2A, and the side of the duplex is leaning onto the surface of the UL52 CTD, specifically at the Zn^2+^-binding end (**Fig. S12C**). An UL5 loop between the *β*25 and α29 forms a pin domain that is positioned at the ssDNA/dsDNA junction and ready to assist the unwinding of dsDNA upon translocation of UL5 (**Fig. S12D**). Together, UL5’s specific binding to the 3’-overhang, the location of dsDNA, and the positioning of the pin domain indicate that UL5 belongs to the SF1A subgroup of SF1 helicases,(*35*) consistent with prior description of HP function.(*19*)

### UL8 auxiliary subunit

Unlike UL5 and UL52 subunits, UL8 has no detectable enzymatic activities. However, it stimulates UL52 primase and UL5 helicase activities in the presence of ICP8.(*36*) UL8 is essential viral DNA replication *in vivo*.(*37*) It interacts with the UL5-UL52 complex, modulates its intracellular localization(*38*), and stimulates the binding of UL5-UL52 to the forked DNA substrate.(*39, 40*) Moreover, UL8 promotes the annealing of complementary ssDNA to form X- and Y-branched duplex DNA structures.(*40*)

Previous homology-based analysis predicted that UL8 shared partial similarity with B-family DNA polymerases and contains palm, fingers, and thumb domains in its C-terminal half.(*41*) Our cryo-EM structure of HP confirmed this prediction and further revealed that the UL8 also contains N-terminal and exonuclease domains resembles closely to a B-family DNA polymerase architecture (**Figs S13 and S14**). However, unlike the representative examples of B-family DNA polymerases, UL8 acquired a more pronounced donut-like shape (**Fig. S13**), that is consistent with the earlier observed ring-shaped structures from negative stain EM(*40*).

For comparative analysis, we selected the well characterized structures of bacteriophage RB69 DNA polymerase (1ig9), and the catalytic core of human DNA polymerase α (Polα) (4qcl).(*42, 43*) RB69 possesses both exonuclease and DNA polymerase activities, while Polα retains only DNA polymerase activity. Their side-by-side comparison with UL8 shows significant topological differences, especially in the N-terminal and exonuclease domains (**Fig. S13A**). Interestingly, the absence of exonuclease activity in Polα was achieved mainly by substitution of active site residues but preserving the overall architecture of the exonuclease fold. In comparison, the exonuclease domain of UL8 exhibits significant topological differences and lacks the bulk of the active site motifs. The loss of UL8 DNA polymerase activity is achieved by substitutions of catalytically active residues without dramatic changes in the folding of conserved regions.(*41*) Inspection of UL8 surface charge also revealed noticeable differences compared with Polα and RB69 DNA polymerase (**Fig. S13B**).

In both latter polymerases, the concentration of positive charge is observed in the middle of proteins comprising the DNA docking areas. However, in the middle of UL8 the positive charge is less prominent, while some cluster is observed in the exonuclease domain (**Fig. S13B**). These comparisons indicate that UL8 is unlikely to bind the template-primer in the same manner as active B-family DNA polymerases. This is consistent with previous DNA-binding studies of UL8 showing that it binds only to ssDNA fragments longer than 50 nucleotides.(*40*)

The presence of an inactivated polymerase-exonuclease module is not unique to the HP complex. Previously, we have discovered two polymerase-exonuclease modules within the catalytic subunit of human DNA polymerase ε (Polε): one is active in the N-terminal part, and another is inactive in the middle of the protein.(*44*) Further cryo-EM studies revealed a role this Polε module played in docking of Polε onto CMG-helicase.(*45, 46*) UL8 may also fulfill a similar role of a structural hub, tethering the components of the HSV1 replisome. Indeed, in addition to its interactions with the UL5-UL52, UL8 has been suggested to interact with other replication factors, including OBP, ICP8, and UL30.(*15, 47, 48*)

Extensive mutagenesis of UL8-charged amino acid residues have revealed several areas apart from the UL52-UL8 interface that might be implicated in interactions with the subunits of HSV1 replisome and/or DNA (**Fig. S14**).(*49*) In particular, the triple mutant A23 (R254A/D255A/D257A) exhibited a lethal phenotype. Double mutant A28 (R378A/E379A) had no effect, while A28 mutations combined with G314E, or with an in-frame eight-amino acid deletion near the N terminus (amino acids 77–84) led to a temperature-sensitive phenotype for DNA replication.(*49*) Moreover, deletion of 26 residues from the N-terminus abolished DNA replication in transient transfection assays(*38*).

In summary, all UL8 mutations that do not disrupt UL5-UL52-UL8 interactions but affect HP activity are mapped to the N-terminal/exonuclease domains (**Figs S14 and S15A**), pointing to them as a potential interaction interface for assembly into the HSV1 replication machinery.

### UL8-UL52 interactions

HP structure shows that the thumb domain of UL8 forms a stable complex with the MD of UL52 (**Fig. 1**). Indeed, the A49 (R640A–D642A–E644A) and A53 (R677A–R678A) mutants of UL8 disrupt the UL8-UL52 interaction and exhibit temperature-sensitive phenotypes.(*49*) In addition, UL52 residues 422–887 are essential for its interaction with UL8.(*20, 50*) Mapping of amino acid residues in the A49 and A53 mutants into the HP structure shows that R640A substitution results in the loss of the inter-subunit salt bridge. However, the remaining substitutions do not participate directly in UL8-UL52 interaction but may reduce the stability of UL8 due to the loss of intra-subunit interactions in the area near the interaction surface. The buried surface area between the UL8 and UL52 is 2,366 Å^2^. This inter-subunit interactions are dominated by hydrophobic contacts (**Fig. S15A**) that are further enhanced by fifteen potential H-bonds: two main-chain-to-main-chain, 10 side-chain-to-main-chain, and three involving side chains only (**Fig. S15B**).

### UL5-UL52 interactions

Previous UL5-UL52 binding studies have mapped the NTD of UL52 as a docking site for UL5.(*20*) However, from our structure, extensive UL5-UL52 interactions are observed in two separate; referred to as the large and small contact areas, with buried surface areas of 5,833 Å^2^ and 1,737 Å^2^, respectively (**Figs S16 and S17**). The large area involves the NTD of UL52, consistent with the binding studies,(*20*) and engages multiple domains of UL5, including the 1A, 1B, 2A, and C-terminal domains, along with only one H-bond involved with domain 2C (**Fig. S16**). The small area involves the CTD of UL52 and the domain 2A of UL5 and has not been previously described (**Fig. S17**).

A hefty amount of H-bonds in the large area consists of one main-chain-to-main-chain type, eighteen side-chain-to-main-chain type, and ten side-chain-to-side-chain type (**Fig. S16A**). Hydrophobic interactions are distributed evenly along the interaction interface (**Fig. S16B**). Interactions in the small area are dominated by H-bonds (**Fig. S17**). Among the fifteen H-bonds, one is of the main-chain-to-main-chain type, six are side-chain-to-main-chain, and eight are between the side chains. Only a few scattered hydrophobic contacts are observed at the UL5-UL52 interface. Among these, the most prominent contacts are formed by P378 of UL5 packed in a small hydrophobic depression on the UL52 CTD surface formed by H948, F949, and F1037 (**Fig. S17**).

### Binding of pritelivir

For the pritelivir-bound HP cryo-EM structure, the map resolution for Lobe A was 3.14 Å (**Figs. S18 and S19**). In this map, the location and conformation of pritelivir were well defined (**Fig. 4A**). The pritelivir fills a partially accessible groove at the UL5-UL52 interface (**Fig. 4B**) and is positioned nearly 12 Å apart from the ATP-binding site of UL52. One end of pritelivir with its 2-pyridyl is wedged deeply into a depression in UL5 located between the loop belonging to the conserved motif IV and the α-helix immediately C-terminal to motif IV (UL5-α_13_) (**Fig. 4C**). The other end of the inhibitor with the aminosulfonyl group sits into the depression between the helix UL52-α_15_ of the helical bundle of UL52 NTD and a bridge helix UL52-α_31_ in the link between the middle and C-terminal domains of UL52 (**Fig. 4C**).

**Figure 4.**
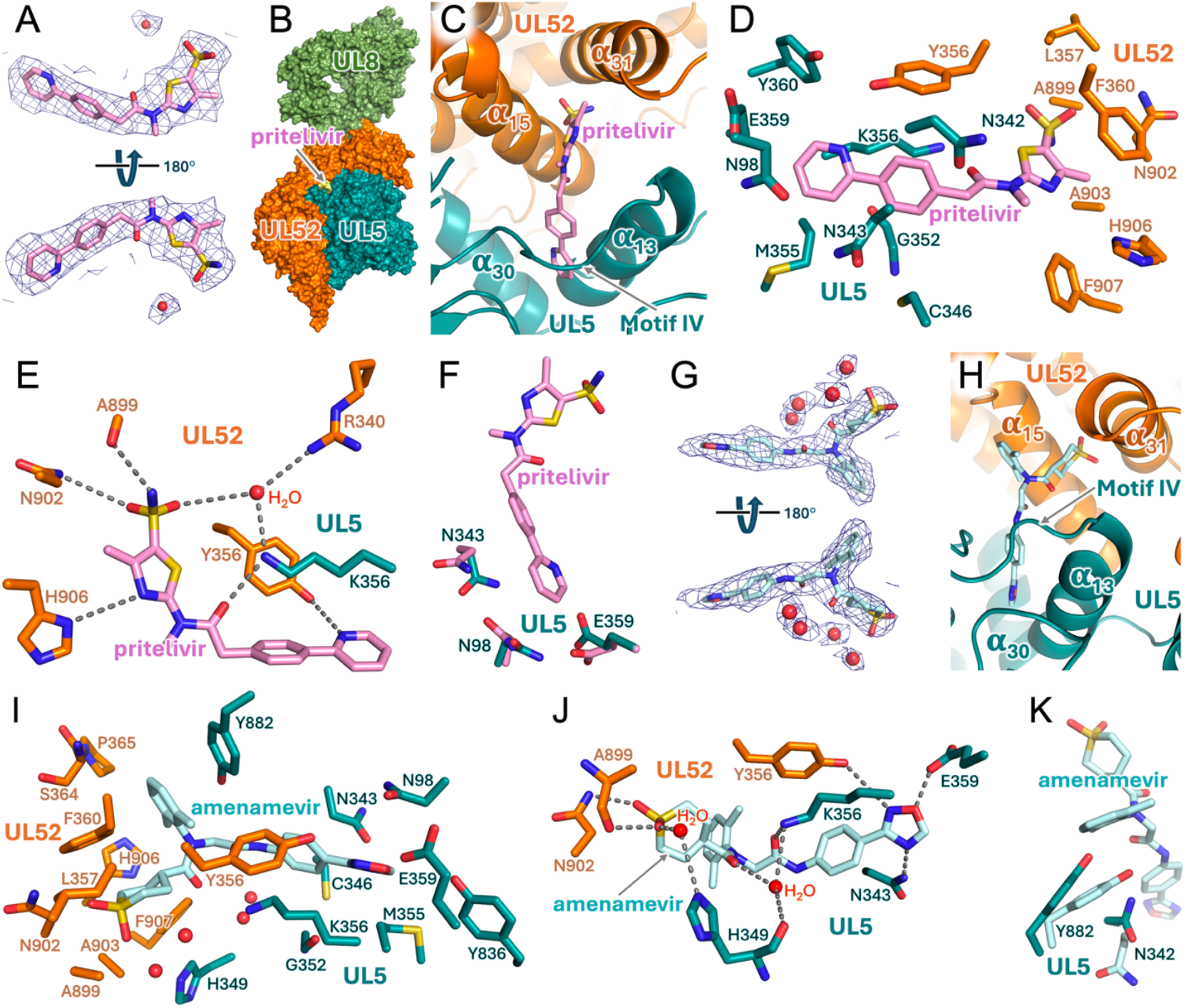
Inhibitor binding by HP-DNA. (**A**) Cryo-EM map of HP-DNA-pritelivir corresponding to a bound pritelivir. (**B**) Surface representation of HP-DNA-pritelivir complex. An arrow is pointed to an area occupied by pritelivir. (**C**) A close-up view of a pritelivir-binding area at the interface of UL5 and UL52. (**D** and **E**) Side chains of UL5 and UL52 amino acid residues forming (**D**) hydrophobic contacts and (**E**) H-bonds with pritelivir. (**F**) Side chains of amino acid residues that change their positions upon binding of pritelivir. (**G**) Cryo-EM map of HP-DNA-amenamevir corresponding to a bound amenamevir. (**H**) A close-up view of an amenamevir-binding area at the interface of UL5 and UL52. (**I** and **J**) Side chains of UL5 and UL52 amino acid residues forming (**I**) hydrophobic contacts and (**J**) H-bonds with amenamevir. (**K**) Side chains of amino acid residues that change their positions upon binding of amenamevir.

Pritelivir forms hydrophobic interactions with the side chains of UL5 amino acid residues N98, N342, N343, C346, M355, K356, E359, Y360, and the α-carbon of G352; and with the side chains of UL52 amino acid residues Y356, L357, F360, A899, N902, A903, H906, and F907 (**Fig. 4D**). In addition, five potential H-bonds were observed between pritelivir and the HP complex. We also observed a density for one well-defined water molecule interacting with both the inhibitor and the protein (**Fig. 4E**). Comparison of the HP-pritelivir structure with the structure of inhibitor-free HP revealed subtle shifts of the residues in and around the binding site. The most notable conformational change occurred in the side-chain conformation of E359 which in turn causes the concerted rearrangement of the side chains of N98 and N343 (**Fig. 4F**).

### Binding of amenamevir

For the amenamevir-bound HP cryo-EM structure, the map resolution for Lobe A was 2.86 Å (**Figs. S20 and S21**). The cryo-EM map has a well-resolved density for amenamevir and the surrounding residues as compared to that of HP-pritelivir complex (**Fig. 4G**). The binding site of amenamevir has a significant overlap with that of pritelivir, with noticeable differences (**Fig. S22A**). Unlike pritelivir with a two-headed elongated structure, amenamevir has a bulkier, three-headed structure (**Fig. S1**). Its azoxime, similar to the 2-pyridyl of pritelivir, is wedged deeply into the depression in UL5 located between the motif IV and the helix UL5-α_13_, while its thiane 1,1-dioxide sits in the depression between the helices UL52-α_15_ and UL52-α_31_, similar to the aminosulfonyl end of pritelivir (**Fig. 4H**). In addition, m-Xylene of amenamevir holds the helix UL52-α_15_ at a surface adjacent to the area covered by its thiane 1,1-dioxide.

The amenamevir-binding surface of HP is formed by the side chains of UL5 amino acid residues N98, N343, C346, H349, M355, K356, E359, Y836, Y882, and the α-carbon of G352; and by the side chains of UL52 amino acid residues Y356, L357, F360, P365, A899, N902, A903, H906, F907 and the α-carbon of S364 (**Fig. 4I**). Six potential H-bonds were observed between amenamevir and the HP complex (**Fig. 4J**). Extra density peaks in proximity to amenamevir were assigned to four water molecules. Two of these water molecules are bridging the amenamevir-protein with H-bonds. The binding of amenamevir causes some changes in the binding area, with the most notable change caused by m-Xylene of amenamevir; it pushes the side chain of UL5 Y882, and that in turn shifts the side chain of UL5 N342 (**Fig. 4K**).

### Pritelivir- and amenamevir-resistant mutations

It is not surprising that all the residues associated with the observed pritelivir-resistant mutations (N342K, G352C, G352R, M355T, K356N, K356T, and K356Q in UL5; and A899T in UL52) are in the list of residues interacting with it. Modeling of these mutations points to a potential disruption of pritelivir docking (**Fig. S22B**). Amenamevir and pritelivir share the positions of some residues with the resistant mutations in UL5 (G352V and M355I), while the positions of the amino acid residues with the amenamevir-resistant mutations in UL52 (S364G and R367H) are different (**Fig. S22C**). Both residues in the helical bundle of the UL52 N-terminal domain are in a proximity of m-Xylene of amenamevir, but over 8 Å away from the pritelivir, explaining the effect of their mutations on amenamevir binding but little or none effect on pritelivir binding.

### Mechanism of HP inhibition

The snapshots of the reaction pathway in SF1 helicases have revealed a nucleotide binding-induced conformational change of the two motor domains.(*33*) Accordingly, UL5 translocation along the DNA also requires significant movements of domains 1A and 2A relative to each other (**Fig. 5A**). They are expected to be closest in ATP/GTP-bound state and furthest apart in the absence of ATP/GTP. All three HP structures presented here do not contain a bound ATP in UL5 active site. Therefore, for comparative analysis, we obtained a model of ATP-bound UL5 using the AlphaFold3 server.(*51*) The predicted model suggests that a potential driving force behind the ATP-facilitated movement of domain 2A is the shift of the conserved R345 and R841 side chains induced by their interactions with the triphosphate moiety of ATP (**Fig. 5B**).

**Figure 5.**
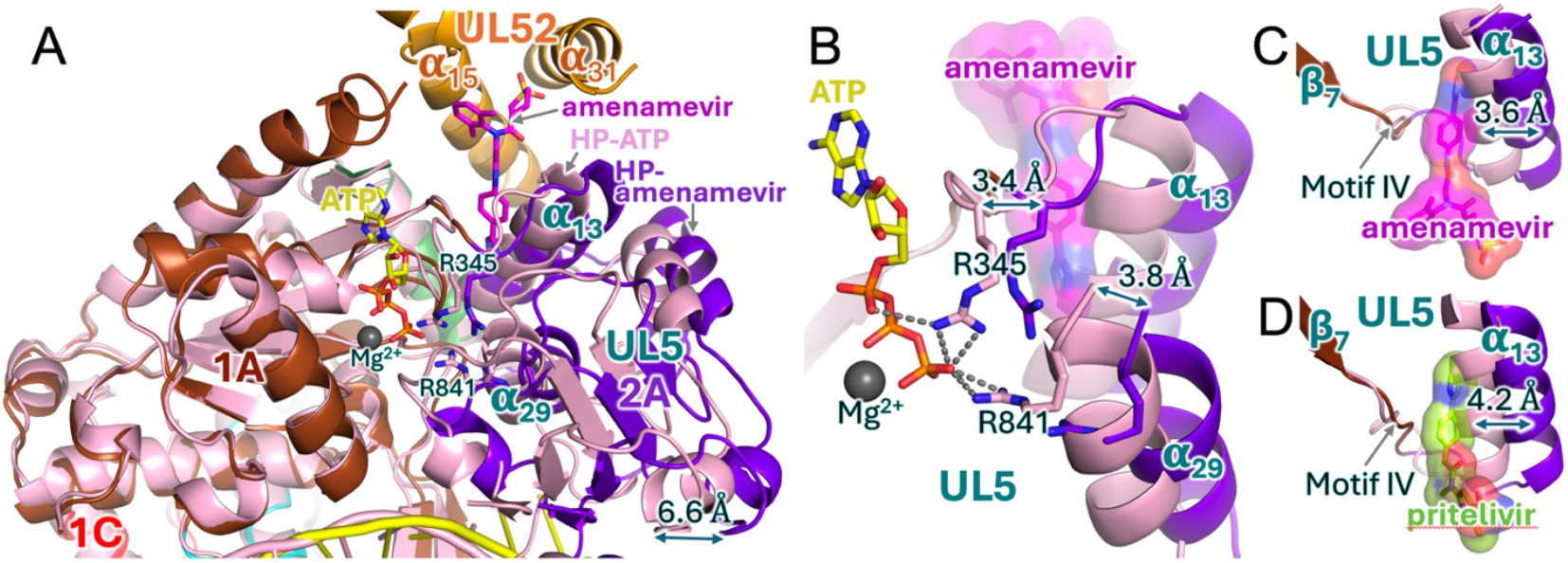
Structural basis of UL5 inhibition by amenamevir and pritelivir. (**A**) Comparison of UL5 structure in a HP-DNA-amenamevir complex determined by cryo-EM and the model of ATP-bound UL5 generated by AlphaFold3.(*51*) To detect the differences caused by ATP binding, we superimposed their 1 A domains. UL5 in cryo-EM structure is colored by domains as in Fig. 3 and UL5-ATP-Mg complex is colored in pink. Comparison shows the shift in domain 2A reaching 6.6 Å. (**B**) A close-up view of the ATP-binding area presented in panel A. The side chains of the conserved arginines R345 and R841 that are responsible for the shift of domain 2A relative to domain 1A are shown as sticks. The shifts of R345 and R841 α-carbons upon ATP binding or dissociation are shown by arrows. Colors are as shown in panel A. (**C**) Amenamevir and (**D**) pritelivir prevent the relative movement of UL5 domains by blocking the movement of UL5 α-helix α_13_.

To estimate the range of the domain movement, we defined the UL5 N-terminal half within domain 1A as a stationary part (because it is anchored to UL52 NTD) while the UL5 C-terminal half within domain 2A as a mobile part (**Fig. 5A**). Superimposition of the domains 1A of ATP-bound UL5 model and UL5 in inhibitor-bound HP cryo-EM structures revealed a significant shift in domain 2A, spanning a 6.6 Å translation in complex with amenamevir (**Fig. 5A**). The helix UL5-α13 of domain 2A, which is adjacent to inhibitor, shifts 3.6 Å and 4.2 Å in complexes with amenamevir and pritelivir, respectively (**Figs 5C and 5D**). These model analyses indicate that sufficient free space between domain 1A (motif IV) and the helix α13 is necessary to accommodate the movement of domain 2A upon ATP/GTP-binding. However, in each inhibitor-bound HP complex, either amenamevir or pritelivir occupies this space (**Figs 5C and 5D**), preventing the relative movement of domains 1A and 2A and thus inhibiting UL5 helicase activity.

## Discussion

Our HP complex cryo-EM structures reveal novel insights into the coordinated action of the helicase and primase subunits at the replication fork. First, HP UL5 binds to a 3’-flap at replication fork and translocates in a direction from 3’ to 5’.(*19*) This indicates that HP-associated herpes polymerase UL30-UL42 translocates together with HP by synthesizing a complimentary strand on an unwound 3’-flap. Second, a 5’-flap of unwound dsDNA is protruding towards the CTD of UL52 and then to the primase active site that is facing the growing 5’-flap. The CTD is expected to be activated by the 5’-flap and the initiating NTP, resulting in its dissociation from UL5 and then positioning the 5’-flap at the primase active site for *de novo* primer synthesis. In other words, the direction of 5’-flap extension is well poised for primers synthesis on the lagging strand (**Fig. S23A**).

Despite different architectures of DNA replication forks in HSV1 and human, they share similarity in how DNA replication is setup. For example, HSV1 UL30-UL42, acting on the 3’-flap extruded from UL5 helicase, resembles the human CMG helicase-bound Polε acting on a leading strand,(*52*) while the 5’-flap in HSV1 HP extends toward the primase active site, similar to the lagging strand extending towards the primase active site in a human CMG helicase-bound Polα-primase.(*53*) Both the human and HSV1 primases lack the initiation of primer synthesis activity in apo forms because the positions of their CTDs are fixed by interactions with Polα(*29, 53*) and UL5 (**Fig. S17**), respectively.

In the human complex, primase becomes active when the length of the 5’-flap reaches 15 to 60 nt (*53*), and the processive synthesis of Okazaki fragments is performed by PCNA-Pol*δ*. In HSV1, this latter function is supposedly fulfilled by UL30-UL42 as suggested in the model of the HSV1 replication fork proposed by the Kuchta and Weller groups.(*18*)

Inhibitor-bound cryo-EM structures reveal the mechanism of HP helicase activity and its modulation by amenamevir and pritelivir. These inhibitors do not compete directly with ATP or GTP for binding to UL5 helicase. Instead, they dock apart from ATP/GTP-binding site in a cavity between the domains 1A and 2A of UL5 and prevent the relative movement of these domains, thus allosterically inhibiting dsDNA unwinding by the helicase (**Fig. S23B**).

While the structure of the key HP complex reported here provides significant insights into the HSV1 replication fork organization, further studies are required to determine the stoichiometry of the replisome components (UL5-UL52-UL8, ICP8, and UL30-UL42) and the structural basis of their concerted actions. Additional efforts are also necessary to establish how the auxiliary HSV1 and the host proteins(*54, 55*) contribute to the complexity of the HSV1 replication fork.

## Author Contributions

A.B. performed functional assays. Q.H. carried out cryo-EM sample vitrification, data collection, and map construction with supervision by C.L. Y.S and L.M.M. carried out cloning, expression, and purification of HP complexes. Y.S. performed protein-DNA binding studies. Y.S. and N.D.B. crystallized HP-DNA and HP-DNA-pritelivir complexes and collected X-ray diffraction data sets. T.H.T. initiated the project, processed X-ray diffraction data, and solved a low-resolution crystal structure of HP-DNA-pritelivir complex, built and refined the high-resolution models of all HP complexes using cryo-EM maps, wrote the manuscript, and prepared the figures with support from A.G.B., Q.H., N.D.B., Y.S. and C.L.

## Data and materials availability

The cryo-EM maps of the HSV1 HP complexes have been deposited to the Electron Microscopy Data Bank (EMDB) under the accession codes EMDB-49585, EMDB-49582, EMDB-49583, and EMDB-49584 for HP-DNA complex, and EMDB-49563, EMDB-49560, EMDB-49561, and EMDB-49562 for HP-DNA-amenamevir complex, and EMDB-49669, EMDB-49586, EMDB-49587, and EMDB-49588 for HP-DNA-pritelivir complex. The corresponding atomic coordinates are deposited in the RSCB Protein Data Bank under accession codes 9nnp (HP-DNA), 9nn2 (HP-DNA-amenamevir), and 9nqp (HP-DNA-pritelivir).

## Supplementary Materials

Materials and Methods

Tables S1 to S4

Figs. S1 to S23

References

