## Supplementary Materials for "Structural Basis of Herpesvirus Helicase-Primase Inhibition by Pritelivir and Amenamevir"

[1]Eppley Institute for Research in Cancer and Allied Diseases, Fred & Pamela Buffett Cancer Center. University of Nebraska Medical Center, Omaha, NE, USA.

[2]Department of Biochemistry, University of Wisconsin-Madison, Madison, WI, USA.

[*]Equal contribution

**The PDF file includes:**

Materials and Methods

Tables S1 to S4

Figs. S1 to S23

References

### Materials and Methods

#### Cloning, expression and purification

cDNAs encoding helicase-primase subunits UL52, UL5, and UL8 of human herpesvirus 1 (strain 17) were obtained from GenScript and individually cloned into the pFastBac1 transfer vector (Invitrogen). Because the project initially was X-ray crystallography-oriented, thirty amino acids predicted to be disordered were deleted from the N-terminus of UL5, and the $His_9$-tag was added to the N-terminus of UL8 followed by a recognition sequence for the precision protease. High-titer viruses were made using Bac-to-Bac Baculovirus Expression System (Invitrogen). $1.8x10^9$ Sf21 cells in 1 L shaking culture were infected simultaneously with recombinant baculoviruses encoding helicase-primase subunits at a multiplicity of infection of 1:2 and were cultivated in an orbital shake (27 ºC, 120 r.p.m.) for 56-60 hours. Cells were harvested by centrifugation at 200 g for 5 min and frozen. Cell pellet was defrosted on ice and lysed in buffer $A_{150}$: 20 mM Tris-HCl, pH 7.9, 150 mM NaCl, 10 mM $K_2HPO_4$, pH 7.9, 2% glycerol, 4 mM β-mercaptoethanol (β-ME), 0.4 mM PMSF, and 1 mg $l^{-1}$ leupeptin. Cell debris was removed by centrifugation at 45,000 g for 15 min, and the obtained lysate was applied to Profinity IMAC resin (7ml, Bio-Rad) charged with $Ni^+$ ions. Column was washed by 70 ml of buffer $A_{150}$, and the complex was eluted by 70 ml gradient of 0.2 M imidazole-HCl, pH 7.8, in buffer $A_{75}$. The peak fractions were combined and loaded to 5 ml Heparin HiTrap HT column (Cytiva), washed by 15 ml buffer $A_{75}$, and eluted by 50 ml gradient of buffer $A_{500}$. The peak fractions were combined, and the His-tag was removed from UL8 during overnight incubation with a precision protease at 4 ºC. The following day, the sample was loaded to a 1-ml monoQ column (Cytiva), washed by 5 ml $bA_{150}$, and eluted by a 20-ml gradient of $bA_{500}$. The peak fractions were combined, dialyzed to 10 mM Hepes-Na, pH 7.5, 150 mM KCl, 1% glycerol, 1 mM DL-dithiothreitol (DTT), concentrated to 2 mg/ml, and frozen. The obtained sample was pure and mainly in a monomeric state, according to dynamic light-scattering analysis (Fig. S12).

#### Preparation of HP complexes

HP without substrates (apo-form, 290 kDa) was concentrated to 5 mg/ml in the buffer containing 20 mM Hepes-NaOH, pH 7.5, 200 mM KCl, 1.5% glycerol, 1 mM DTT. HP in complex with DNA and adenosine 5′-(β,γ-imido)triphosphate (AMP-PNP, a non-hydrolysable analogue of

ATP) was prepared by adding a ten base-pair DNA duplex containing one and fifteen nucleotides long 5'- and 3'-flaps, respectively, which was made by annealing oligonucleotides 11a and 25b (**Table S1**). The complex was concentrated to 6 mg/ml in the buffer containing 10 mM Hepes-Na, pH 7.5, 140 mM KCl, 1% glycerol, 4 mM DTT, 1.6 mM $MgCl_2$, and 1 mM AMP-PNP. In HP complexes with DNA, AMP-PNP, and an inhibitor (100 μM pritelivir or 40 μM amenamevir), DNA contains a 27 bp duplex and 16 nucleotides long 5'- and 3'-flaps (obtained by annealing oligonucleotides 43a and 43b). These complexes were concentrated to 6-7 mg/ml in the buffer containing 15 mM Hepes-Na, pH 7.5, 150 mM KCl, 1% glycerol, 3 mM DTT, 1.6 mM $MgCl_2$, and 1 mM AMP-PNP. All complexes were aliquoted by 5μl and flash-frozen in liquid nitrogen.

**Cryo-EM sample grids preparation and data collection**

The frozen samples were thawed immediately before cryo-EM grid preparation. The holey electron microscopy grids (C-Flat 1.2/1.3, 300 mesh) were glow-discharged for 20 seconds at 15 mA (PELCO easiGlow) before use. A final concentration of 2 mM CHAPSO(*56*) was added to the UL5-UL8-UL52-DNA-pritelivir complex samples and UL5-UL8-UL52-DNA-Amenamevir complex samples, whereas 4 mM CHAPSO was added to the UL5-UL8-UL52-DNA complex samples. Subsequently, 3.5 μl of the prepared samples was deposited onto the freshly glow-discharged grids. The grids were blotted for 4 s at 4 °C and 95% relative humidity before being plunge-frozen in liquid ethane using a Vitrobot Mark IV (Thermo Fisher).

The sample grids were initially screened for particle density and ice quality on an Arctica 200 kV transmission electron microscope (Thermo Fisher) equipped with a BioQuantum K3 energy filter detector (Gatan). The grids with the best quality were selected for data collection using a Titan Krios transmission electron microscope (Thermo Fisher) at the National Cryo-Electron Microscopy Facility at the National Cancer Institute of the National Institutes of Health. The UL5-UL8-UL52-DNA-Amenamevir complex sample dataset (12,660 movies) was collected using the Latitude software at a calibrated pix size of 1.07 Å per pixel, a defocus range from −1.0 to 2.5 μm, a total dose of 50.0 $e^{-}$ $Å^{-2}$ (40 frames), and the K3 detector (20-eV energy slit) in counting mode. The UL5-UL8-UL52-DNA-pritelivir complex sample dataset (9,216 movies) was collected using Latitude software at a calibrated pix size of 1.11 Å per pixel, a defocus range from −1.0 to −2.5 μm, a total dose of 50.0 $e^{-}$ $Å^{-2}$ (40 frames), and the K3 detector (20-eV energy slit) in counting mode. The UL5-UL8-UL52-DNA complex sample dataset (12,553 movies) was collected using the Latitude software at a calibrated pix size of 1.07 Å per pixel, a defocus range from −1.0 to −2.5 μm, a total dose of 50.0 $e^{-}$ $Å^{-2}$ (40 frames), and the K3 detector (20-eV energy slit) in counting mode.

**Cryo-EM data processing**

In general, all three datasets were processed using a similar image processing strategy. The movie frames were aligned, and their constant transfer function (CTF) parameters were calculated using CryoSPARC's patch motion correction and patch CTF estimation.(*57*)

For the UL5-UL8-UL52-DNA-Amenamevir complex sample dataset, a total of 11,242 micrographs remained after curation based on particle density and ice quality. Particles picked from 500 representative micrographs using the template-free blob-picker were extracted and subjected to two rounds of 2D classification to remove most junk particles. The best 2D classes were selected as the template for the template picker in CryoSPARC, which was used for the subsequent automated picking for all micrographs. A total of 3,799,736 particles were picked and extracted from all micrographs at 4x binning (4.28 Å per pixel). The particles were subjected to two rounds of 2D classification to remove junk particles, after which the selected particles (750,198 particles) were sorted into three reference-free 3D classes using ab initio modeling in CryoSPARC. These three 3D classes were used as templates for two rounds of heterogeneous refinement, after which the selected particles (904,751) were re-extracted at the original pixel size (1.07 Å per pixel). These particles were subjected into one round of heterogeneous refinement to remove poor-quality particles. A total of 725,979 particles were selected for the subsequent 3D reconstruction using non-uniform refinement in CryoSPARC.(*58*) The particles were further refined with one round of global CTF refinement and reference-based motion correction in CryoSPARC, culminating in non-uniform refinement to a global resolution (reported at Fourier shell correlation of 0.143) of 3.10 Å.

For the UL5-UL8-UL52-DNA-pritelivir complex sample dataset, a total of 8,820 micrographs remained after curation based on particle density and ice quality. Particles picked from 500 representative micrographs using the template-free blob-picker were extracted and subjected to two rounds of 2D classification to remove most junk particles. The best 2D classes were selected as the template for the template picker in CryoSPARC, which was used for the subsequent automated picking for all micrographs. A total of 3,516,384 particles were picked and extracted from all micrographs at 4x binning (4.44 Å per pixel). The particles were subjected to a single round of 2D classification to remove junk particles, after which the selected particles (729,585 particles) were sorted into three reference-free 3D classes using ab initio modeling in CryoSPARC. The particles from two classes (609,139 particles) were re-extracted at the original pixel size before performing another two rounds of ab initio modeling to remove poor-quality particles. A total of 409,183 suitable particles were selected for the 3D reconstruction using non-uniform refinement in CryoSPARC.(*58*) The particles were further refined with one round of global CTF refinement, reference-based motion correction, and local CTF refinement in CryoSPARC, culminating in non-uniform refinement to a global resolution (reported at Fourier shell correlation of 0.143) of 3.42 Å.

For the UL5-UL8-UL52-DNA complex sample dataset, a total of 11,628 micrographs remained after curation based on particle density and ice quality. Particles picked from 500 representative micrographs using the template-free blob-picker were extracted and subjected to two rounds of 2D classification to remove most junk particles. The best 2D classes were selected as the template for the template picker in CryoSPARC for the subsequent automated picking for all micrographs. A total of 4,446,980 particles were picked and extracted from all micrographs at 4x binning (4.28 Å per pixel). The particles were subjected to a single round of 2D classification to remove junk particles, after which the selected particles (906,191 particles) were sorted into three reference-free 3D classes using ab initio modeling in CryoSPARC. The particles from one class (569,272 particles) were re-extracted at the original pixel size before performing another two rounds of ab initio modeling to remove poor-quality particles. A total of 432,244 suitable particles were selected for the 3D reconstruction using non-uniform refinement in CryoSPARC and further refined with one round of global CTF refinement and reference-based motion correction in CryoSPARC. Non-uniform refinement results in a global resolution (reported at Fourier shell correlation of 0.143) of 3.43 Å.

The three-dimensional (3D) variable analysis (3DVA) principal component analysis was performed for three datasets in CryoSPARC and revealed two large swiveling motions of UL5-$UL52_N$-$UL52_C$ domain and UL8-$UL52_{aa410\text{-}893}$ domain. To further improve the quality of the consensus maps, local refinement was performed individually for the UL5-$UL52_N$-$UL52_C$ domain and UL8-$UL52_{aa410\text{-}893}$ domain. The docked models were used as segmentation guides to generate the masks of both domains in ChimeraX software.(*59*) Each domain was subjected to particle subtraction and local refinement in CryoSPARC to yield a final local refined domain EM density map. The two local refined domain EM density maps were combined using the consensus map as a guide. Local resolution evaluations were determined using the local resolution estimation in CryoSPARC with a refined half-map and visualized using ChimeraX software.

**Model building, refinement and validation**

The initial model of HP complex in apo-form was generated by web-based AlphaFold.(*51*) Then each domain from the model was docked into the cryo-EM map of the complex without inhibitors using ChimeraX software(*59*). The obtained structure was inspected with respect to its fit in the cryo-EM map and manually corrected using Coot(*60*). Then, the well-defined portion of ssDNA and two zinc ions were modeled into the map using Coot. The obtained structure was then subjected to real-space refinement using the Phenix software package.(*61*) This structure without an inhibitor was used as a starting model for the HP complexes with inhibitors, amenamevir and pritelivir. The coordinates and restraints files for the inhibitors were generated using the eLBOW module in Phenix. Each inhibitor was placed manually into the corresponding map, and the structures of the complexes with amenamevir and pritelivir were adjusted and refined similar to the HP complex without an inhibitor. The cryo-EM map to model validation statistics was derived from Phenix's comprehensive validation package.

Cryo-EM data collection, refinement, and validation statistics for HP-DNA, HP-DNA-pritelivir and HP-DNA-amenamevir complexes are summarized in **Tables S2-S4.**

**Table S1**. **Oligonucleotides used in this study**

| Name | Sequence (5' → 3' direction; complementary sequences underlined) | Study |
|---|---|---|
| 11a | TGGTGCCGTGG | cryo-EM |
| 25b | CCACGGCACCTACATAATACATACA | |
| 43a | ACATACATAGTACTTAAGGACGAGCTGCAGACGTCCAGGCTGC | |
| 43b | GCAGCCTGGACGTCTGCAGCTCGTCCTATCTGTTCACCTGTCA | |
| 30a | ACATACATAGTACTTGCAGGTCAGTCGAGC | EMSA |
| 30b | GCTCGACTGACCTGCTAACTGATACATACA | |
| 15a | GCAGGTCAGTCGAGC | |
| 15b | GCTCGACTGACCTGC | |

**Table S2**. **Cryo-EM data collection, refinement, and validation statistics for HP-DNA complex**

| | **HP-DNA-complex Composite map** (EMDB-49585) (PDB 9nnp) | **HP-DNA-complex** (consensus map) (EMDB-49582) | **HP-DNA-complex** (UL5-UL52$_N$-UL52$_c$-ssDNA-focused map) (EMDB-49583) | **HP-DNA-complex** (UL8-UL52$_{aa410-893}$-focused map) (EMDB-49584) |
|---|---|---|---|---|
| **Data collection and processing** | | | | |
| Magnification | 81,000x | 81,000x | 81,000x | 81,000x |
| Voltage (kV) | 300 | 300 | 300 | 300 |
| Electron exposure (e–/Å$^2$) | 50.0 | 50.0 | 50.0 | 50.0 |
| Defocus range (μm) | −1.0 to −2.5 | −1.0 to −2.5 | −1.0 to −2.5 | −1.0 to −2.5 |
| Pixel size (Å) | 1.07 | 1.07 | 1.07 | 1.07 |
| Symmetry imposed | C1 | C1 | C1 | C1 |
| Initial particle images (no.) | 4,446,980 | 3,799,736 | 3,799,736 | 3,799,736 |
| Final particle images (no.) | 431,843 | 431,843 | 431,843 | 431,843 |
| Map resolution (Å) | 3.2 | 3.43 | 3.16 | 3.18 |
| FSC threshold | 0.143 | 0.143 | 0.143 | 0.143 |
| **Refinement** | | | | |
| Initial model used (PDB code) | AlphaFold | | | |
| Model resolution (Å) | 3.4 | | | |
| FSC threshold | 0.5 | | | |
| Model resolution range (Å) | 3.1-3.4 | | | |
| Map sharpening *B* factor (Å$^2$) | - | | | |
| Q-Score | 0.51 | | | |
| Model composition | | | | |
| Non-hydrogen atoms | 19498 | | | |
| Protein residues | 2510 | | | |
| Ligands | 2 | | | |
| *B* factors (Å$^2$) | | | | |
| Protein | 69.28 | | | |
| Nucleotide | 63.19 | | | |
| Ligand | 134.19 | | | |
| R.m.s. deviations | | | | |
| Bond lengths (Å) | 0.002 (0) | | | |
| Bond angles (°) | 0.545 (2) | | | |
| Validation | | | | |
| MolProbity score | 1.86 | | | |
| Clashscore | 6.14 | | | |
| Poor rotamers (%) | 1.70 | | | |
| Ramachandran plot | | | | |
| Favored (%) | 95.00 | | | |
| Allowed (%) | 4.72 | | | |
| Disallowed (%) | 0.28 | | | |

**Table S3. Cryo-EM data collection, refinement, and validation statistics for HP-DNA-pritelivir complex**

| | **HP-DNA-pritelivir complex Composite map** (EMDB-49669) (PDB 9nqp) | **HP-DNA-pritelivir complex** (consensus map) (EMDB-49586) | **HP-DNA-pritelivir complex** (UL5-UL52$_N$-UL52$_c$-ssDNA-focused map) (EMDB-49587) | **HP-DNA-pritelivir complex** (UL8-UL52$_{aa410-893}$-focused map) (EMDB-49588) |
|---|---|---|---|---|
| **Data collection and processing** | | | | |
| Magnification | 81,000x | 81,000x | 81,000x | 81,000x |
| Voltage (kV) | 300 | 300 | 300 | 300 |
| Electron exposure (e–/Å$^2$) | 50.0 | 50.0 | 50.0 | 50.0 |
| Defocus range (μm) | −1.0 to −2.5 | −1.0 to −2.5 | −1.0 to −2.5 | −1.0 to −2.5 |
| Pixel size (Å) | 1.11 | 1.11 | 1.11 | 1.11 |
| Symmetry imposed | C1 | C1 | C1 | C1 |
| Initial particle images (no.) | 3,516,384 | 3,516,384 | 3,516,384 | 3,516,384 |
| Final particle images (no.) | 409,125 | 409,125 | 409,125 | 409,125 |
| Map resolution (Å) | 3.1 | 3.42 | 3.14 | 3.20 |
| FSC threshold | 0.143 | 0.143 | 0.143 | 0.143 |
| **Refinement** | | | | |
| Initial model used (PDB code) | AlphaFold | | | |
| Model resolution (Å) | 3.3 | | | |
| FSC threshold | 0.5 | | | |
| Model resolution range (Å) | 3.1-3.4 | | | |
| Map sharpening *B* factor (Å$^2$) | - | | | |
| Q-Score | 0.516 | | | |
| Model composition | | | | |
| Non-hydrogen atoms | 19558 | | | |
| Protein residues | 2512 | | | |
| Ligands | 3 | | | |
| *B* factors (Å$^2$) | | | | |
| Protein | 65.93 | | | |
| Nucleotide | 54.26 | | | |
| Ligand | 39.68 | | | |
| R.m.s. deviations | | | | |
| Bond lengths (Å) | 0.003 (0) | | | |
| Bond angles (°) | 0.521 (0) | | | |
| Validation | | | | |
| MolProbity score | 2.05 | | | |
| Clashscore | 10.14 | | | |
| Poor rotamers (%) | 1.3 | | | |
| Ramachandran plot | | | | |
| Favored (%) | 93.21 | | | |
| Allowed (%) | 6.43 | | | |
| Disallowed (%) | 0.36 | | | |

**Table S4**. **Cryo-EM data collection, refinement, and validation statistics for HP-DNA-amenamevir complex**

| | **HP-DNA-amenamevir complex Composite map** (EMDB-49563) (PDB 9nn2) | **HP-DNA-amenamevir complex** (consensus map) (EMDB-49560) | **HP-DNA-amenamevir complex** (UL5-UL52$_{N}$-UL52$_{c}$-ssDNA-focused map) (EMDB-49561) | **HP-DNA-amenamevir complex** (UL8-UL52$_{aa410-893}$-focused map) (EMDB-49562) |
|---|---|---|---|---|
| **Data collection and processing** | | | | |
| Magnification | 81,000x | 81,000x | 81,000x | 81,000x |
| Voltage (kV) | 300 | 300 | 300 | 300 |
| Electron exposure (e–/Å$^2$) | 50.0 | 50.0 | 50.0 | 50.0 |
| Defocus range (μm) | −1.0 to −2.5 | −1.0 to −2.5 | −1.0 to −2.5 | −1.0 to −2.5 |
| Pixel size (Å) | 1.07 | 1.07 | 1.07 | 1.07 |
| Symmetry imposed | C1 | C1 | C1 | C1 |
| Initial particle images (no.) | 3,799,736 | 3,799,736 | 3,799,736 | 3,799,736 |
| Final particle images (no.) | 725,691 | 725,691 | 725,691 | 725,691 |
| Map resolution (Å) | 2.90 | 3.10 | 2.86 | 2.96 |
| FSC threshold | 0.143 | 0.143 | 0.143 | 0.143 |
| **Refinement** | | | | |
| Initial model used (PDB code) | AlphaFold | | | |
| Model resolution (Å) | 3.1 | | | |
| FSC threshold | 0.5 | | | |
| Model resolution range (Å) | 2.8-3.1 | | | |
| Map sharpening *B* factor (Å$^2$) | - | | | |
| Q-Score | 0.545 | | | |
| Model composition | | | | |
| Non-hydrogen atoms | 19563 | | | |
| Protein residues | 2510 | | | |
| Ligands | 3 | | | |
| *B* factors (Å$^2$) | | | | |
| Protein | 68.03 | | | |
| Nucleotide | 44.08 | | | |
| Ligand | 37.56 | | | |
| R.m.s. deviations | | | | |
| Bond lengths (Å) | 0.003 (0) | | | |
| Bond angles (°) | 0.529 (3) | | | |
| Validation | | | | |
| MolProbity score | 2.4 | | | |
| Clashscore | 17.65 | | | |
| Poor rotamers (%) | 2.55 | | | |
| Ramachandran plot | | | | |
| Favored (%) | 94.96 | | | |
| Allowed (%) | 4.76 | | | |
| Disallowed (%) | 0.28 | | | |

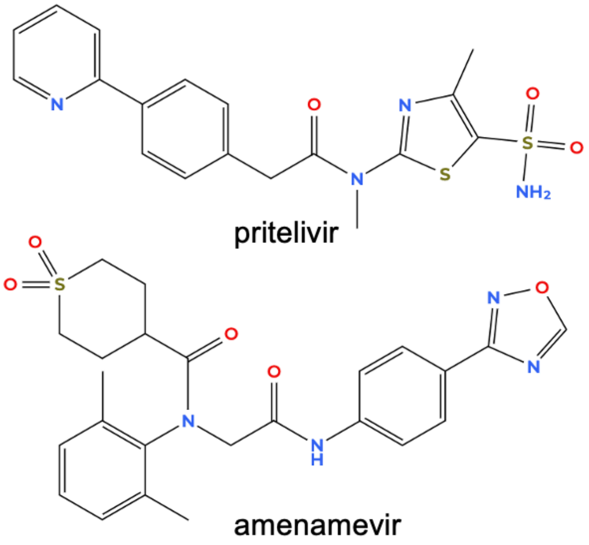

**Figure S1. Helicase-primase inhibitors.**

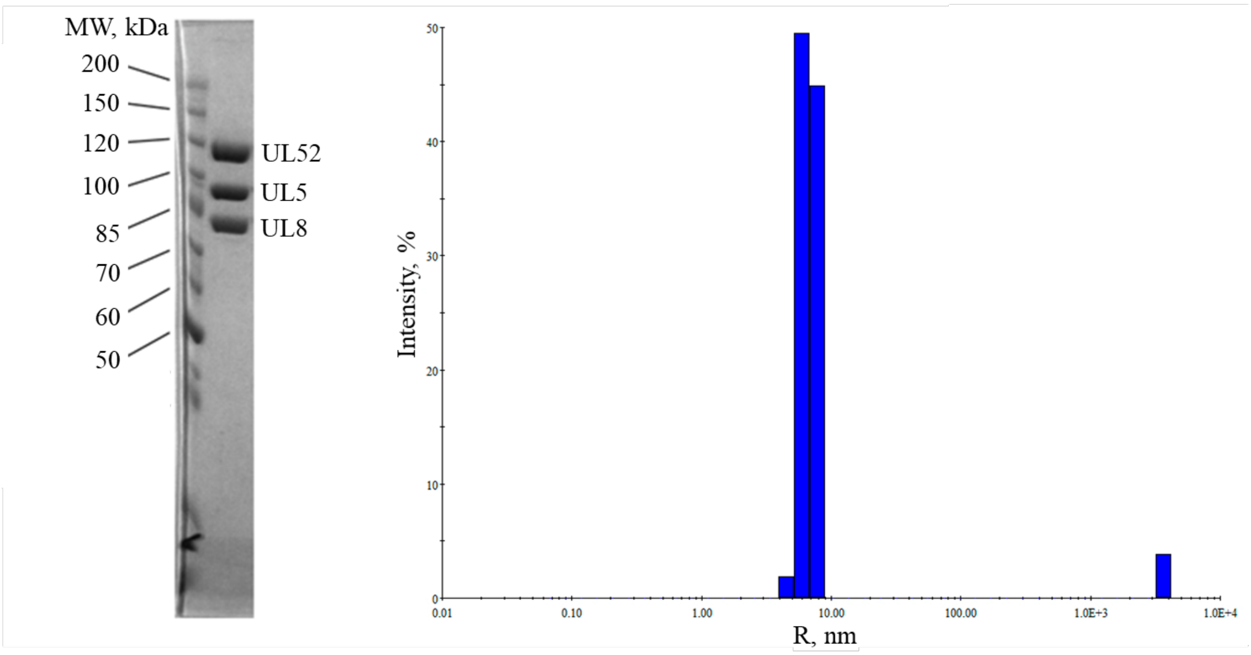

**Figure S2**. **Analysis of HP sample quality. (A)** HP subunits were separated by electrophoresis in 8% SDS-PAGE and stained by Coomassie R-250. Molecular weight markers are shown on the left. **(B)** Results of dynamic light-scattering analysis of HP sample at the concentration of 2 mg/ml. The main peak corresponds to particles with a molecular weight of 299 kDa (hydrodynamic radius is 6.8 nm). Analysis was performed on DynaPro Nanostar (Wyatt technology).

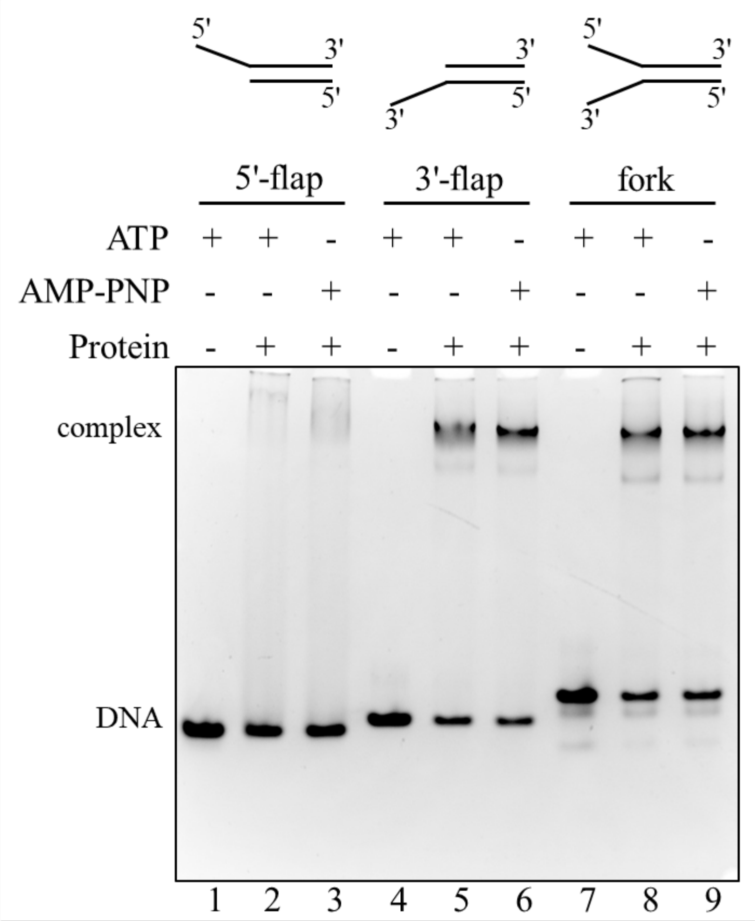

**Figure S3**. **3'-flap is critical for HP binding to the forked DNA substrate.** Reactions containing 3.5 µM HP, 3.5 µM DNA, 2 mM $MgCl_2$, and 0.5 mM AMP-PNP, were incubated for 60 min at 37 ºC. The products were resolved by 5% native PAGE and stained by ethidium bromide. DNA substrates with a 5'-flap, a 3'- flap, and both flaps, were prepared by annealing the oligonucleotides 30a:15b, 30b:15a, and 30a:30b, respectively.

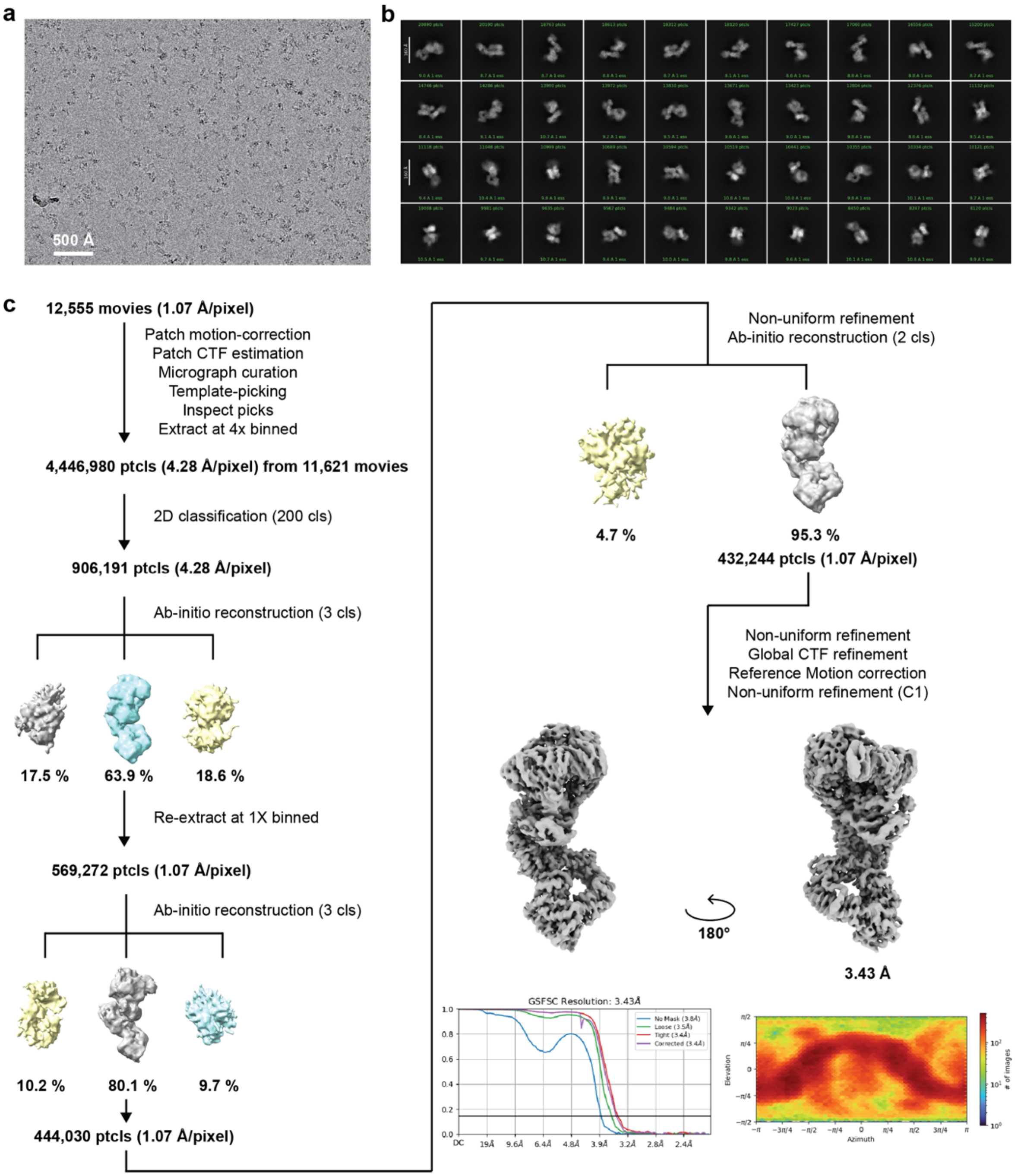

**Figure S4. Cryo-EM processing pipeline for the UL5-UL8-UL52-DNA complex**. (**a**) Representative motion-corrected micrograph (n=12,555) of the cryo-EM dataset for processing. The scale bar dimension is 500 Å. (**b**) 2D classification averages of the UL5-UL8-UL52-DNA complex. A 170 Å scale bar is shown as a white bar. (**c**) The cryo-EM dataset processing pipeline to obtain the consensus map of the UL5-UL8-UL52-DNA complex.

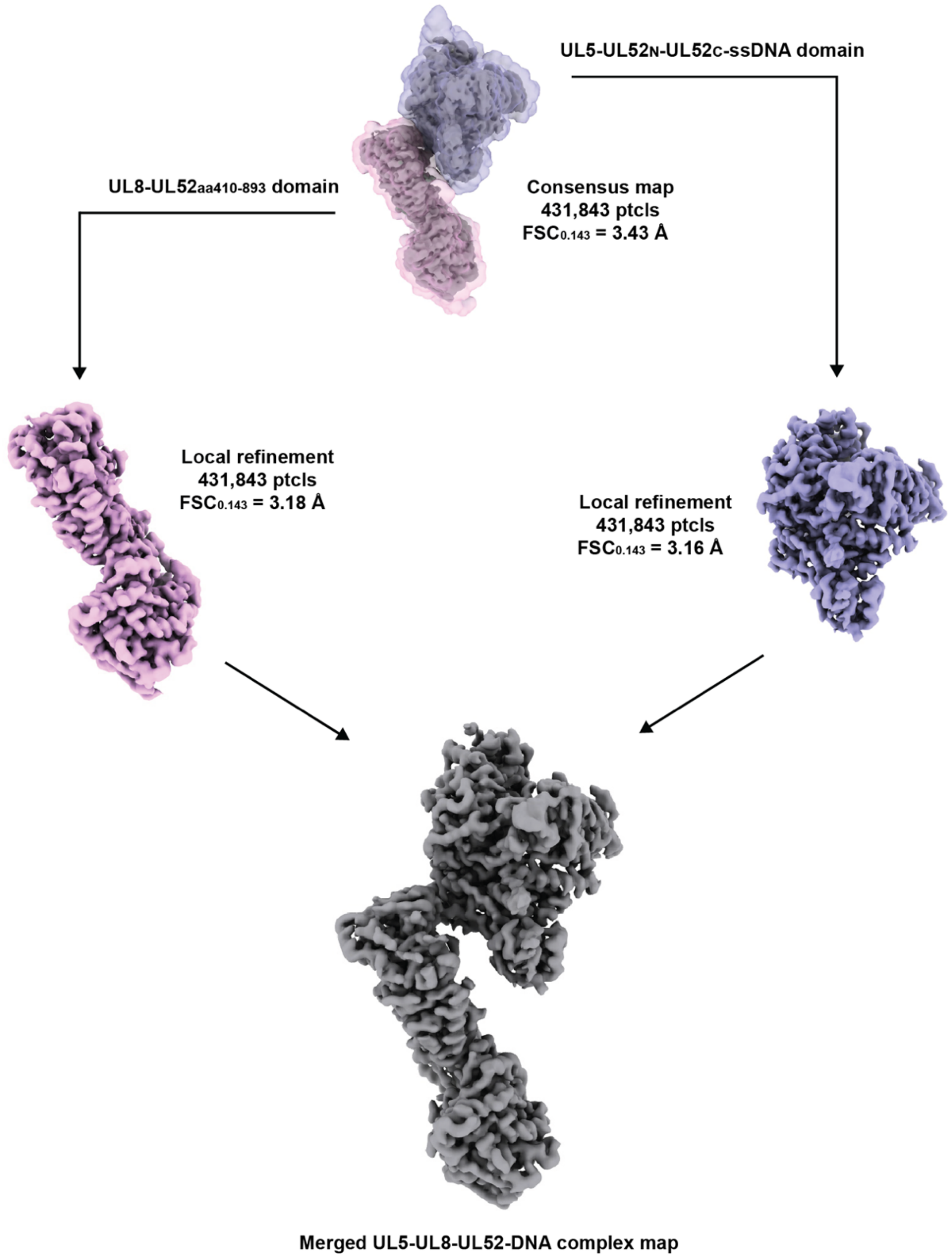

**Figure S5. Local refinement of individual flexible domains of the UL5-UL8-UL52-DNA complex**. The UL5-UL8-UL52-DNA complex was segmented into two distinct domains for particle subtraction and 3D reconstruction. The local masks are shown as semi-transparent. The two domains are merged to yield a better-quality assembled complex map.

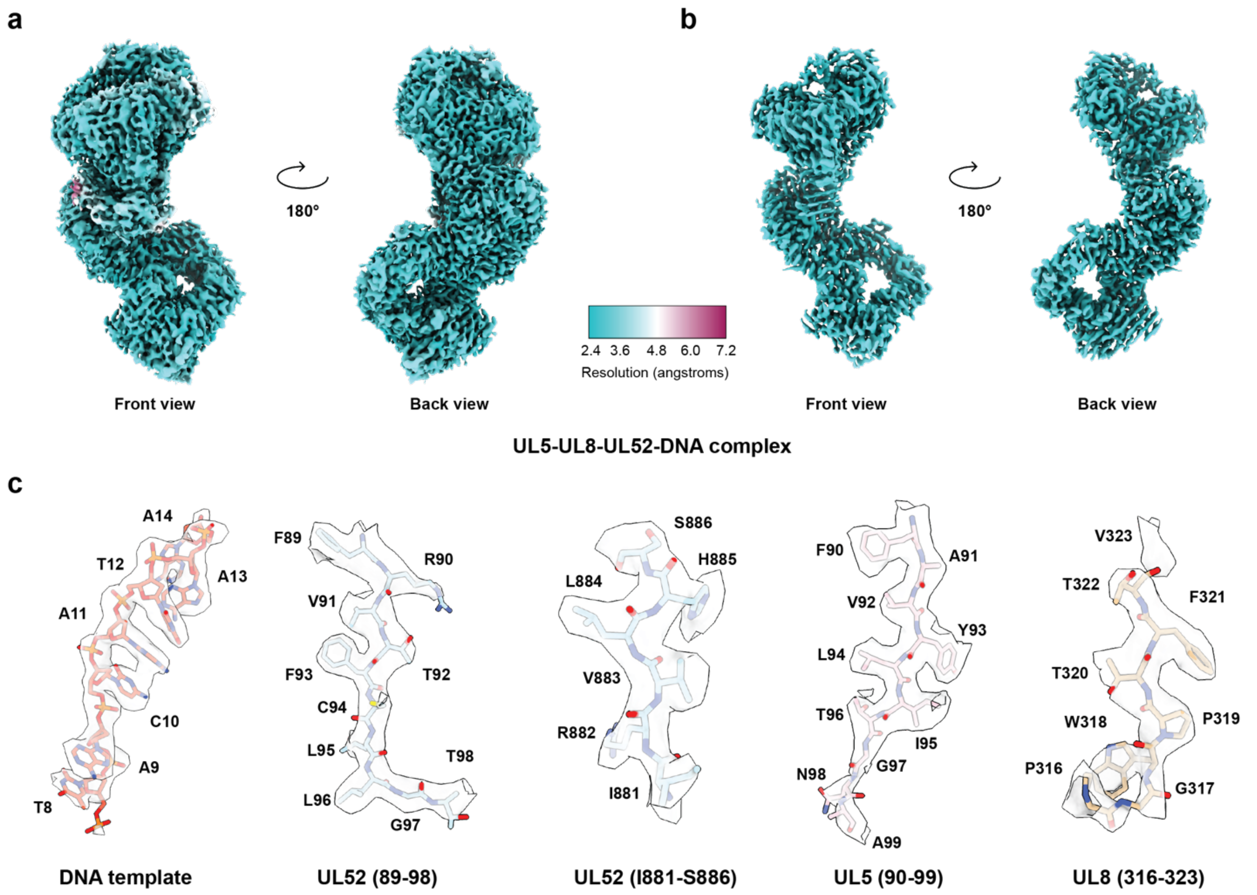

**Figure S6. Local resolution maps and map to model comparison of the cryo-EM reconstruction of the UL5-UL8-UL52-DNA complex**. (**a**) Local resolution maps of the UL5-UL8-UL52-DNA complex at volume threshold of 0.06. (**b**) The same as A but maps are shown at volume threshold of 0.13 to show the higher-resolution regions. **(c)** Representative cryo-EM densities encasing the related models of the UL5-UL8-UL52-DNA complex.

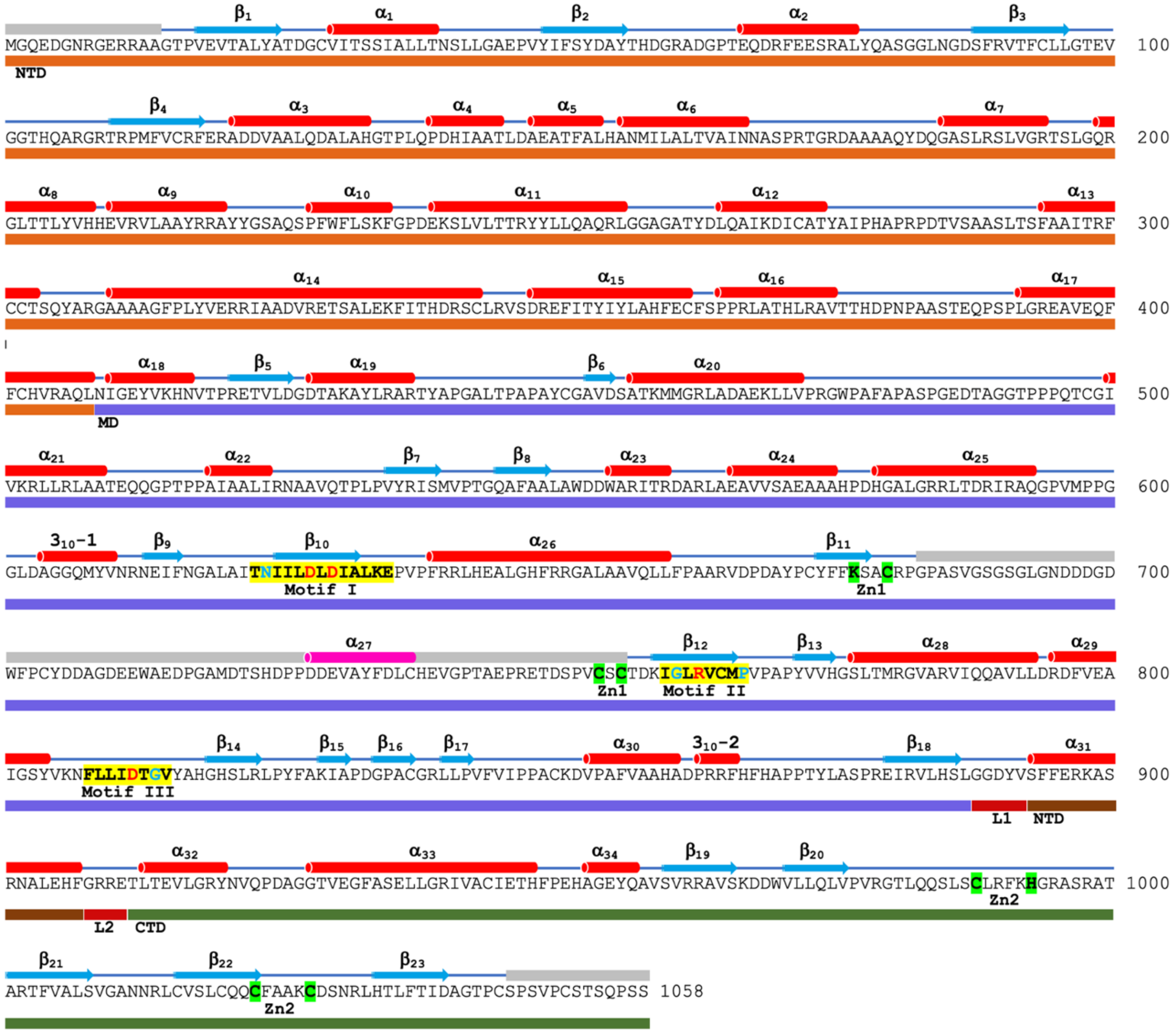

**Figure S7. Amino acid sequence of HSV1 UL52.** Secondary structure elements are shown above the sequences. The α- and $3_{10}$-helices are shown as cylinders in red, β-strands are shown as arrows in cyan, and disordered regions are shown as grey bars. The bars below the sequences correspond to UL52 domains: NTD in orange, MD in slate, CTD in split pea green, and linkers L1 and L2 in firebrick. Zn1 and Zn2 coordinating residues are highlighted by green marker and AEP signature motifs I-III are highlighted by yellow marker. The AEP motif I-III residues conserved in all primases are colored in red and within herpesviruses in cyan.

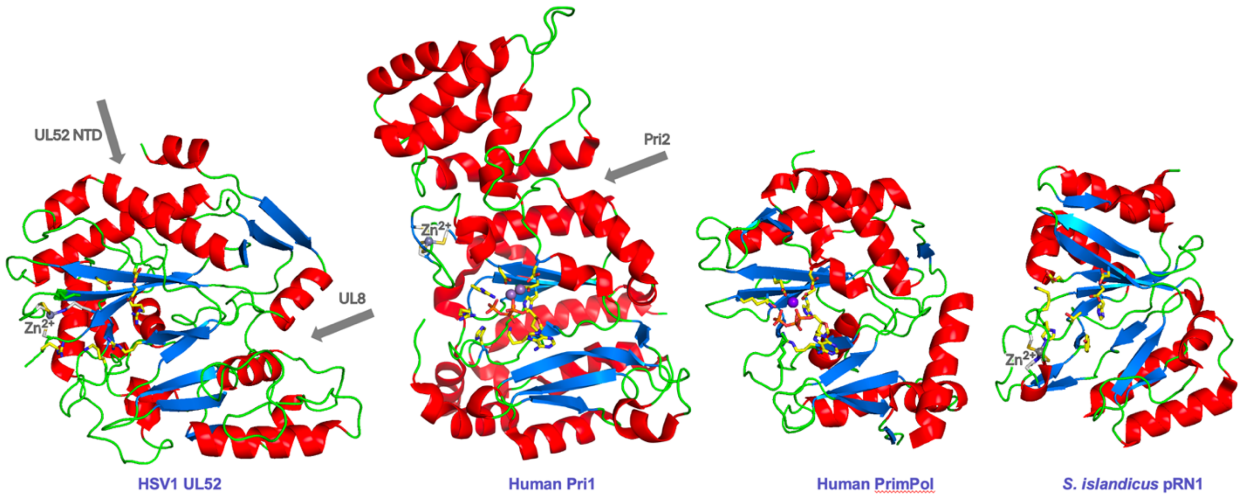

**Figure S8. Side-by-side comparison of primase domains from HSV1 UL52, human Pri1, human PrimPol, and *S. islandicus* pRN1.** The α- and $3_{10}$-helices are shown in red, β-strands in cyan, and turns or random coils in green. The $Zn^{2+}$ and $Mg^{2+}$ ions are shown as grey and purple spheres. The side chains of NTP- and metal-binding residues are shown as sticks. The arrows indicate the locations of UL52 NTD and UL8 relative to primase catalytic domain in HSV1 UL52, and Pri2 relative to Pri1 in human primase. The figures were produced using the coordinates with the following PDB codes: 9nnp for HSV1 HP (current work), 6r5d for human Pri1,(*26*) 5l2x for human PrimPol,(*24*) and 3m1m for *S. islandicus* pRN1.(*25*)

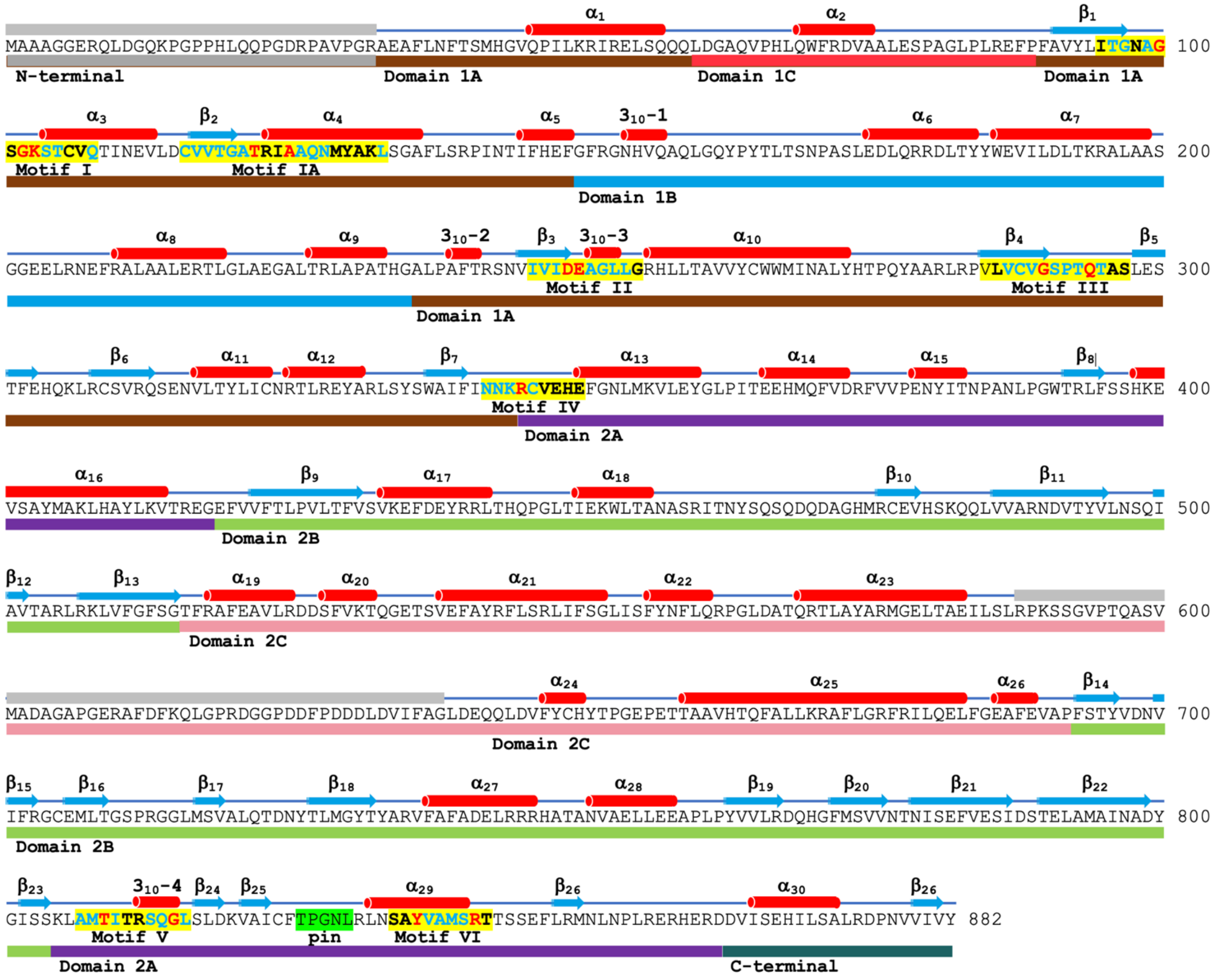

**Figure S9. Amino acid sequence of HSV1 UL5 subunit.** Secondary structure elements are shown above the sequences. The α- and $3_{10}$-helices are shown as cylinders in red, β-strands are shown as arrows in cyan, and disordered regions are shown as grey bars. The bars below the sequences correspond to UL5 domains: N-terminal in grey, 1A in brown, 1B in cyan, 1C in red, 2A in purple, 2B in pale green, 2C in pink, and C-terminal in forest green. The pin domain residues are highlighted by green marker, and the residues of the helicase conserved motifs I-VI are highlighted by yellow marker. The motifs I-VI residues conserved in SF1 helicases are colored in red and within herpesviruses in cyan.

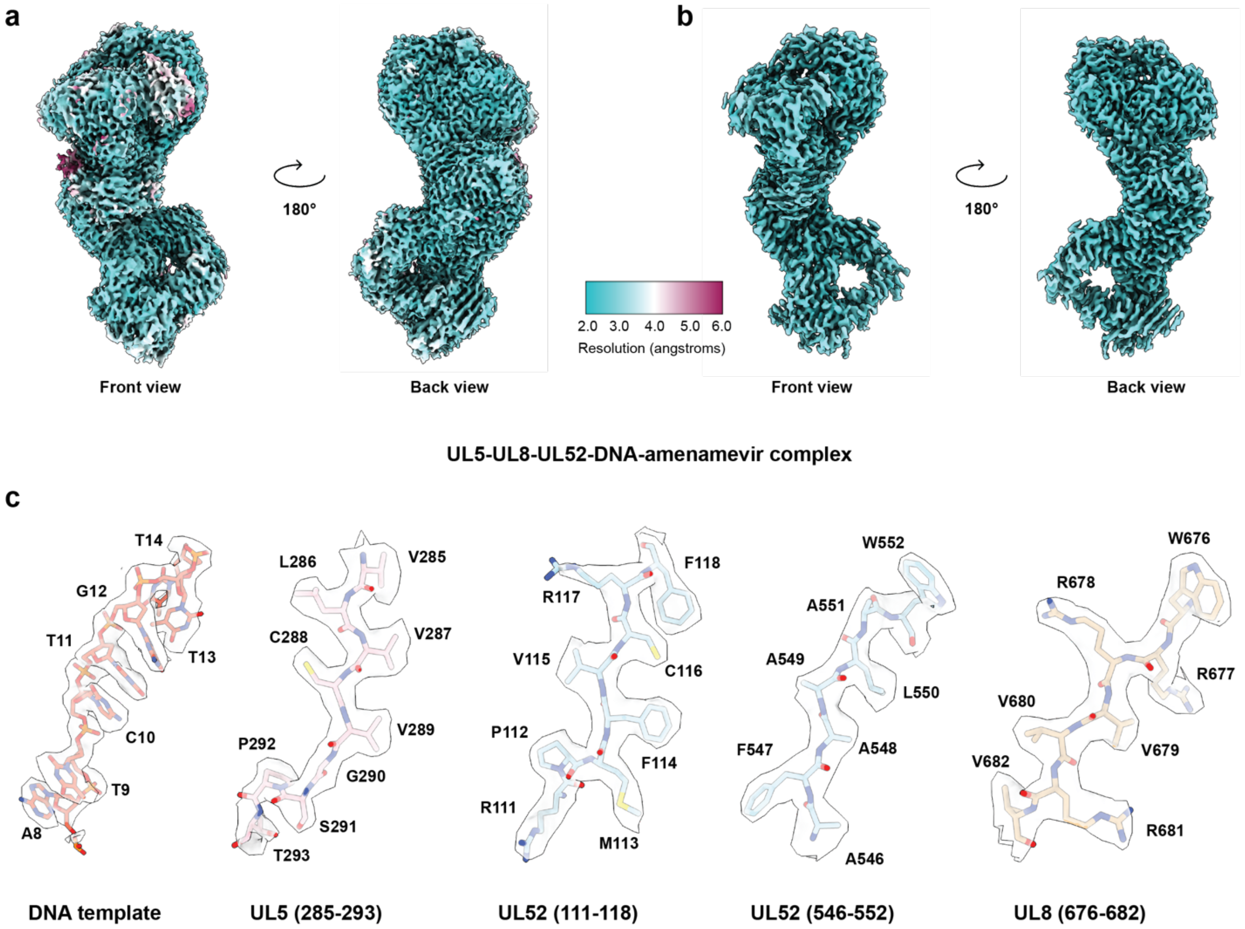

**Figure S10. Local resolution maps and map to model comparison of the cryo-EM reconstruction of** the UL5-UL8-UL52-DNA-amenamevir complex. (**a**) Local resolution maps of the UL5-UL8-UL52-DNA-amenamevir complex at volume threshold of 0.06. (**b**) The same as A but maps are shown at volume threshold of 0.13 to show the higher-resolution regions. **(c)** Representative cryo-EM densities encasing the related models of the UL5-UL8-UL52-DNA-amenamevir complex.

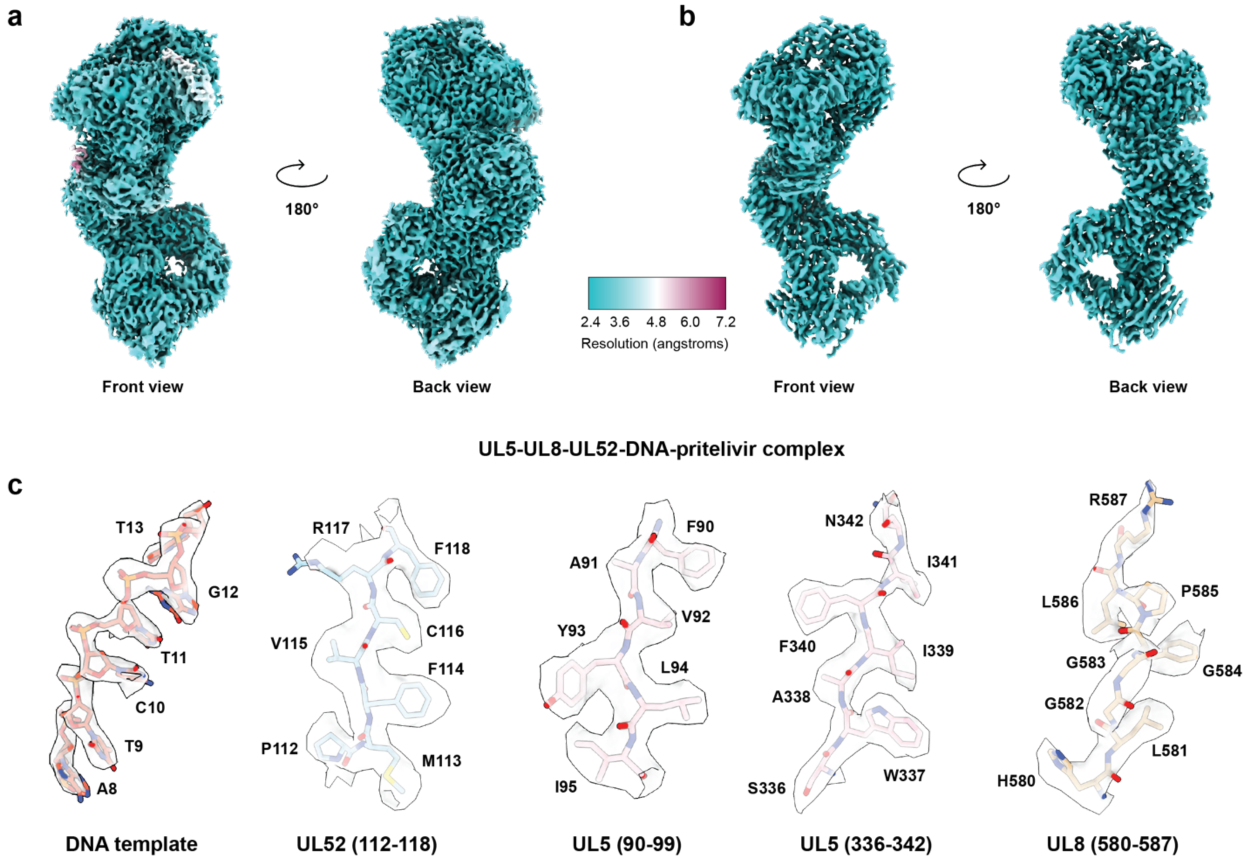

**Figure S11. Local resolution maps and map to model comparison of the cryo-EM reconstruction of the UL5-UL8-UL52-DNA-pritelivir complex**. (**a**) Local resolution maps of the UL5-UL8-UL52-DNA-pritelivir complex at volume threshold of 0.06. (**b**) The same as A but maps are shown at volume threshold of 0.13 to show the higher-resolution regions. **(c)** Representative cryo-EM densities encasing the related models of the UL5-UL8-UL52-DNA-pritelivir complex.

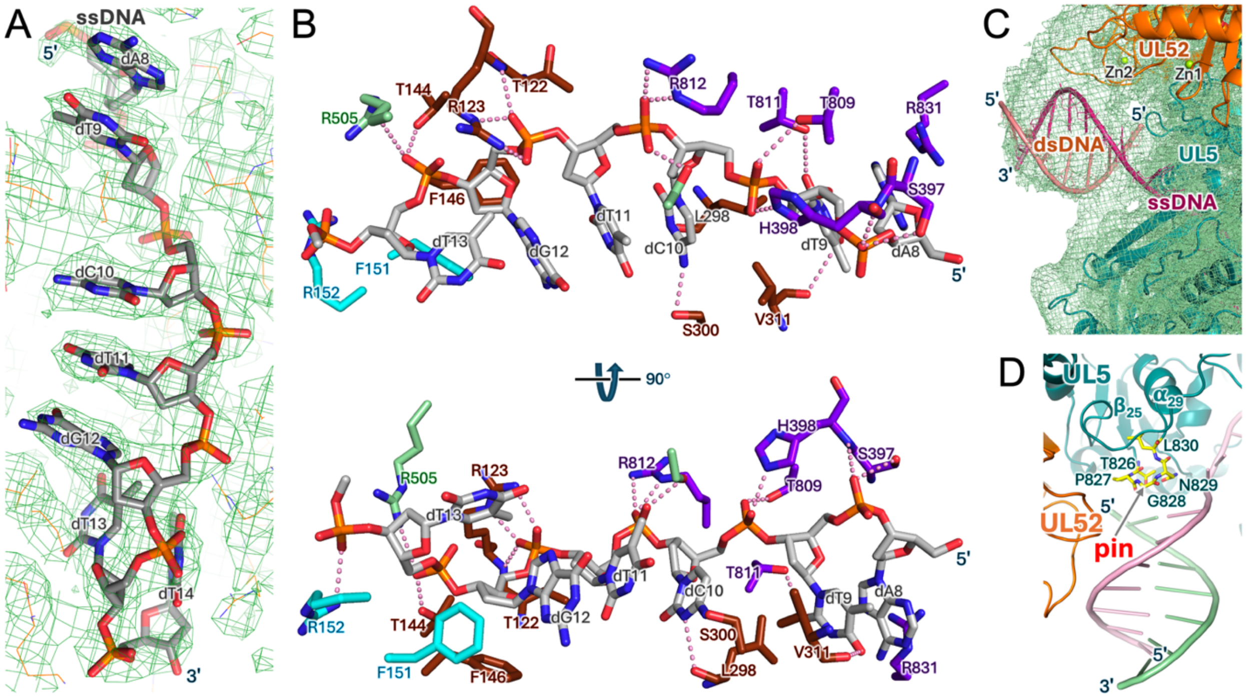

**Figure S12**. **DNA binding by UL5.** (**A**) ssDNA bound fitted into the cryo-EM map. (**B**) H-bonds between UL5 residues and ssDNA. H-Bonds are drawn by dashed lines. For clarity, only interacting amino acid residues are shown and are colored by domain colors as in Figure 2. Two views are shown. (**C**) Unsharpened cryo-EM map reveals the density for dsDNA. (**D**) Location of UL5 pin domain (colored in yellow) relative to DNA.

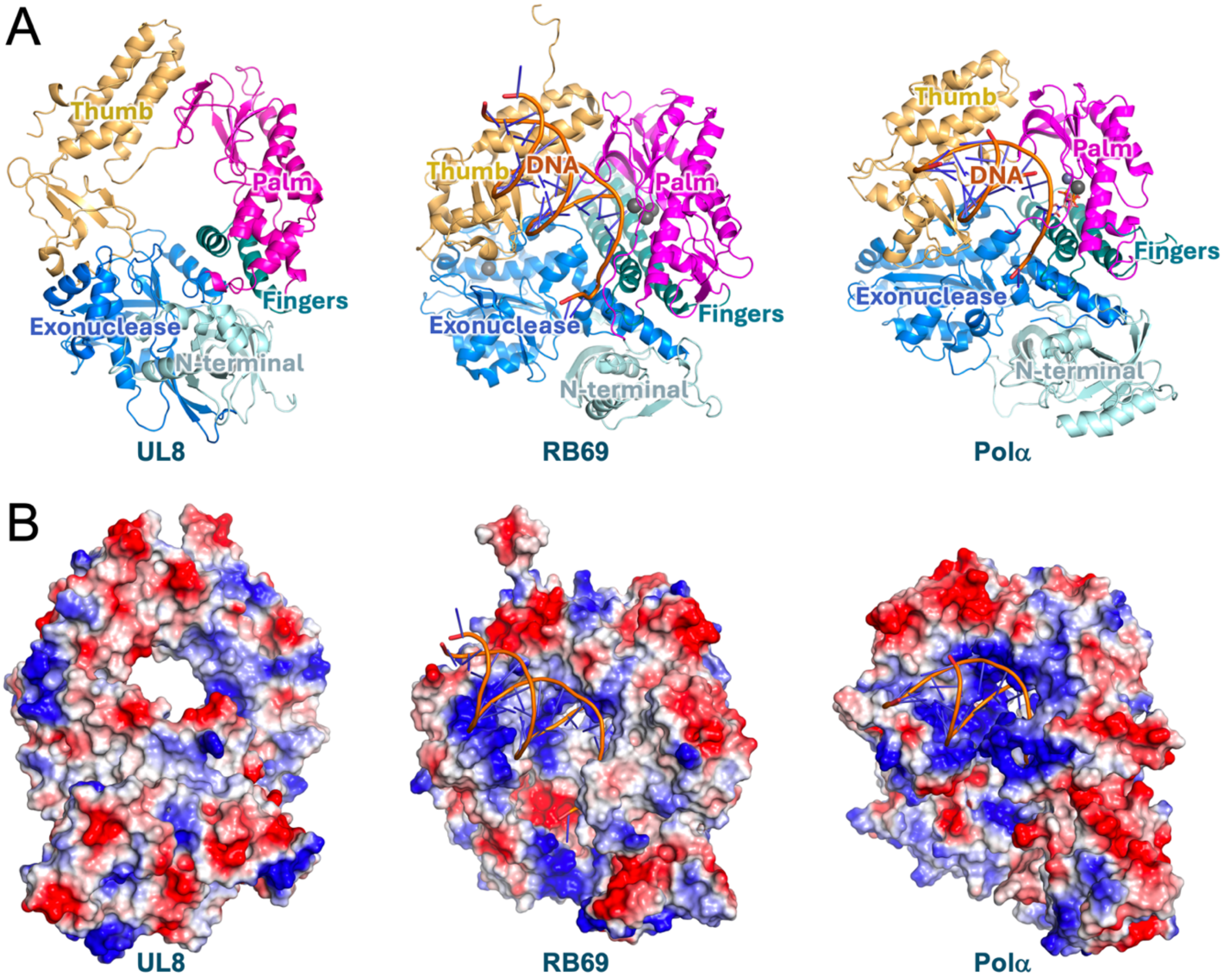

**Figure S13. Structure of HSV1 HP UL8 subunit.** (**A**) Cartoon representation of UL8 and its side-by-side comparison with the bacteriophage RB69 DNA polymerase (PDB code 1ig9) and the human Polα catalytic cores PDB code (PDB code 4qcl).(*42, 43*) The structures are colored by domains: N-terminal in cyan, Exonuclease in blue, Palm in magenta, Fingers in deep teal, and Thumb in light orange. (**B**) Charge distribution on surfaces of UL8, RB69, and Polα. Positively charged areas are in blue and negatively charged areas are in red.

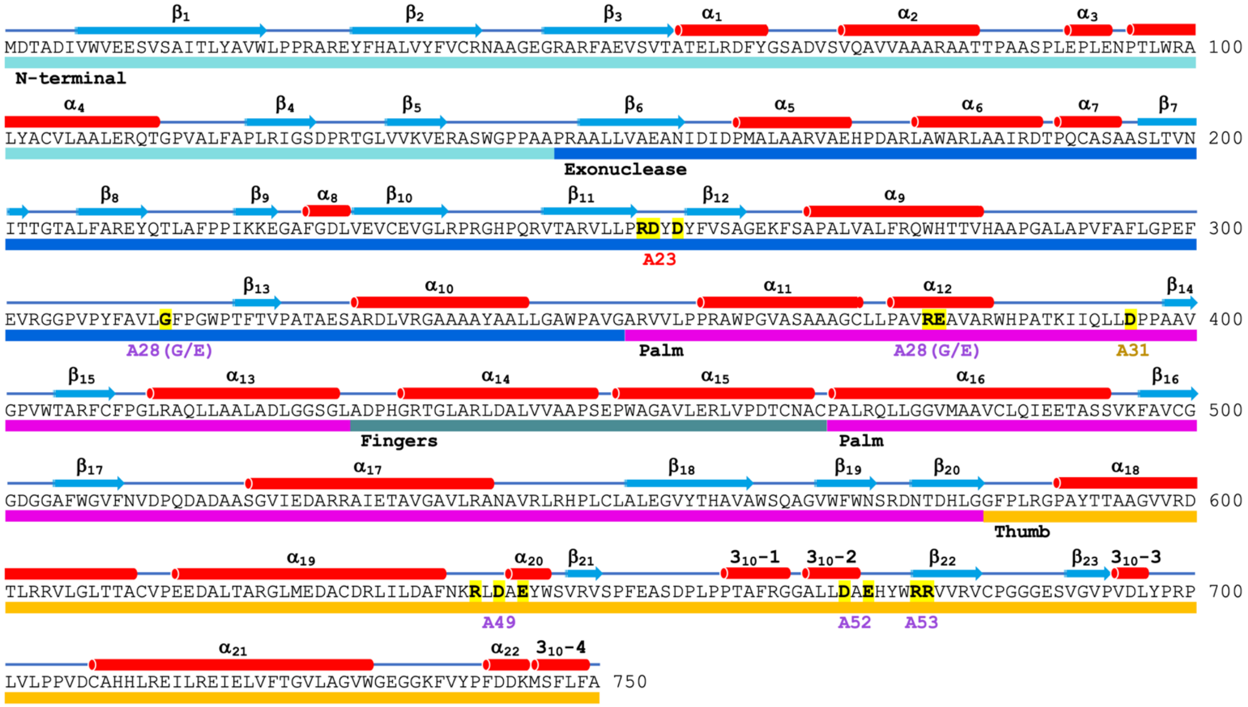

**Figure S14. Amino acid sequence of HSV1 UL8 subunit.** Secondary structure elements are shown above the sequences and labeled. The α- and $3_{10}$-helices are shown as cylinders in red, β-strands are shown as arrows in cyan. The bars below the sequences correspond to UL8 domains: N-terminal in cyan, Exonuclease in blue, Palm in magenta, Fingers in deep teal, and Thumb in light orange. The amino acid residues that were subjected to mutational studies are highlighted by yellow marker. The mutant exhibiting a lethal phenotype is in red, while the mutants having temperature-sensitive phenotypes are in magenta.

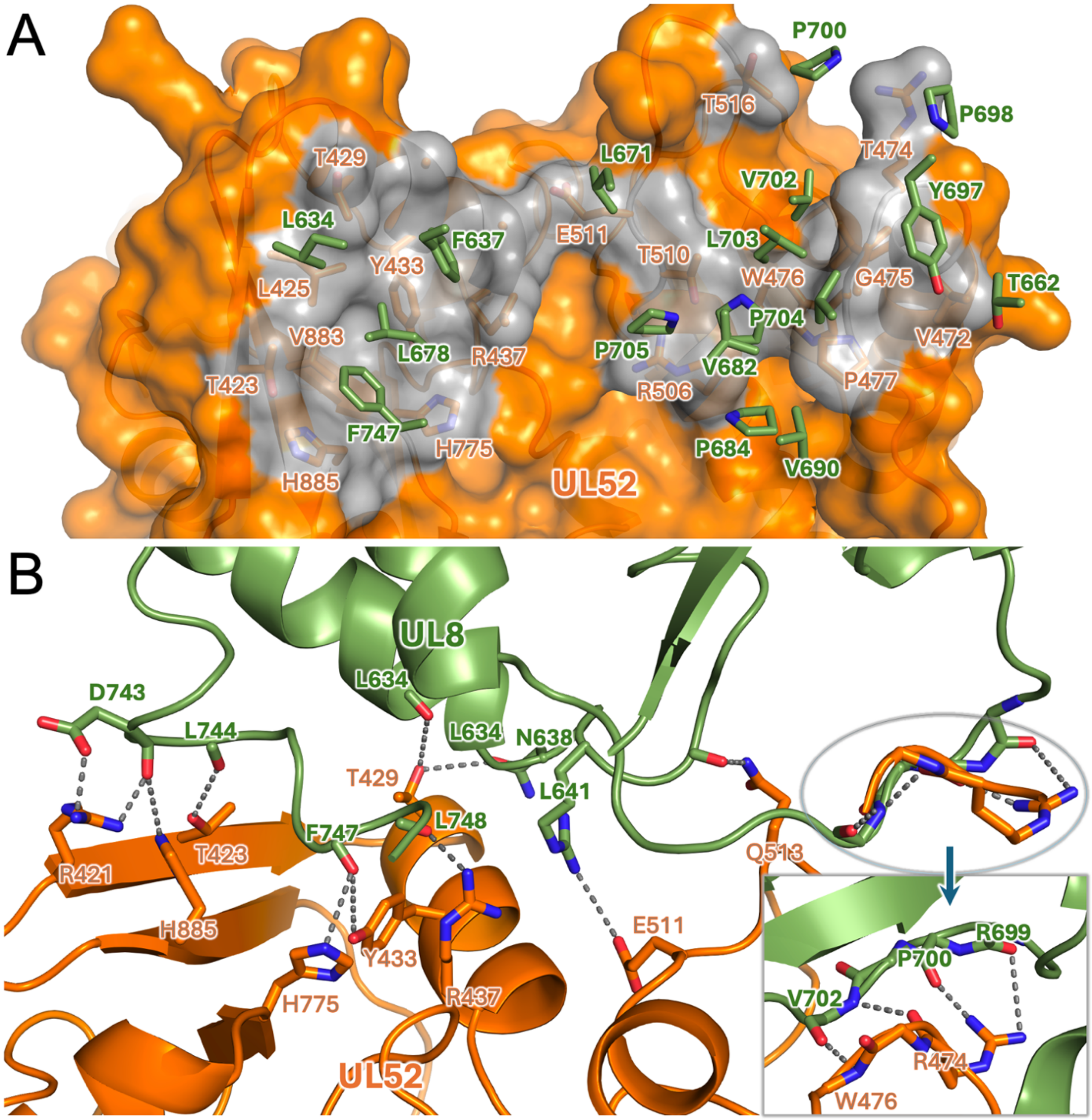

**Figure S15. UL8-UL52 interactions.** (**A**) Hydrophobic interactions between UL8 and UL52. The semitransparent surface of UL8 contributing to hydrophobic interactions is highlighted in grey. The participating residues are shown as sticks and labeled. For the clarity, the cartoon of UL8 is not displayed. (**B**) H-bonds between UL8 and UL52. The H-bonds are drawn as dashed lines. The participating residues are shown as sticks and labeled.

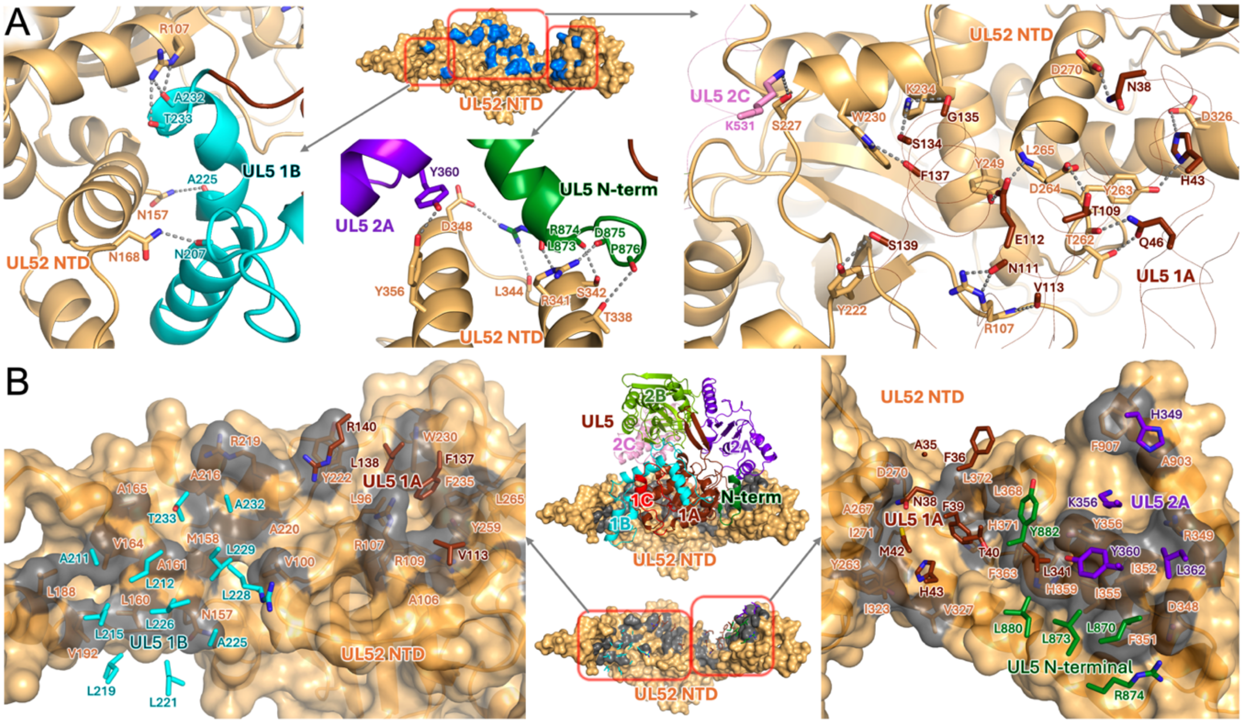

**Figure S16. Interactions between UL5 and NTD of UL52.** (**A**) H-bonds between UL5 and NTD of UL52. Surface representation of the NTD of UL52 is shown in the middle. The surface areas colored in blue correspond to the residues of UL5 NTD participating in H-bond formations with UL5. Three separate images show H-bonds in the enclosed areas in detail. The H-bonds are drawn as dashed lines. The participating residues are shown as sticks and labeled. UL5 domains are color coded as in Figure 3. The UL5 fold on the right image is displayed with a thin line for an unobstructed view of interactions. (**B**) Hydrophobic interactions between UL5 and UL52. The middle-upper image shows the NTD of UL52 in a surface representation and UL5 in a cartoon representation. The latter is color coded by domains. The middle lower image shows the UL8 NTD surface with the grey-highlighted areas contributing to hydrophobic interactions. The detailed hydrophobic interactions in the two enclosed areas are shown on the left and right images. In these images, the semitransparent surface of UL52 NTD contributing to hydrophobic interactions is highlighted in grey. The residues participating in inter-subunit hydrophobic interactions are shown as sticks and labeled. UL5 residues are color coded by domains as in Figure 3.

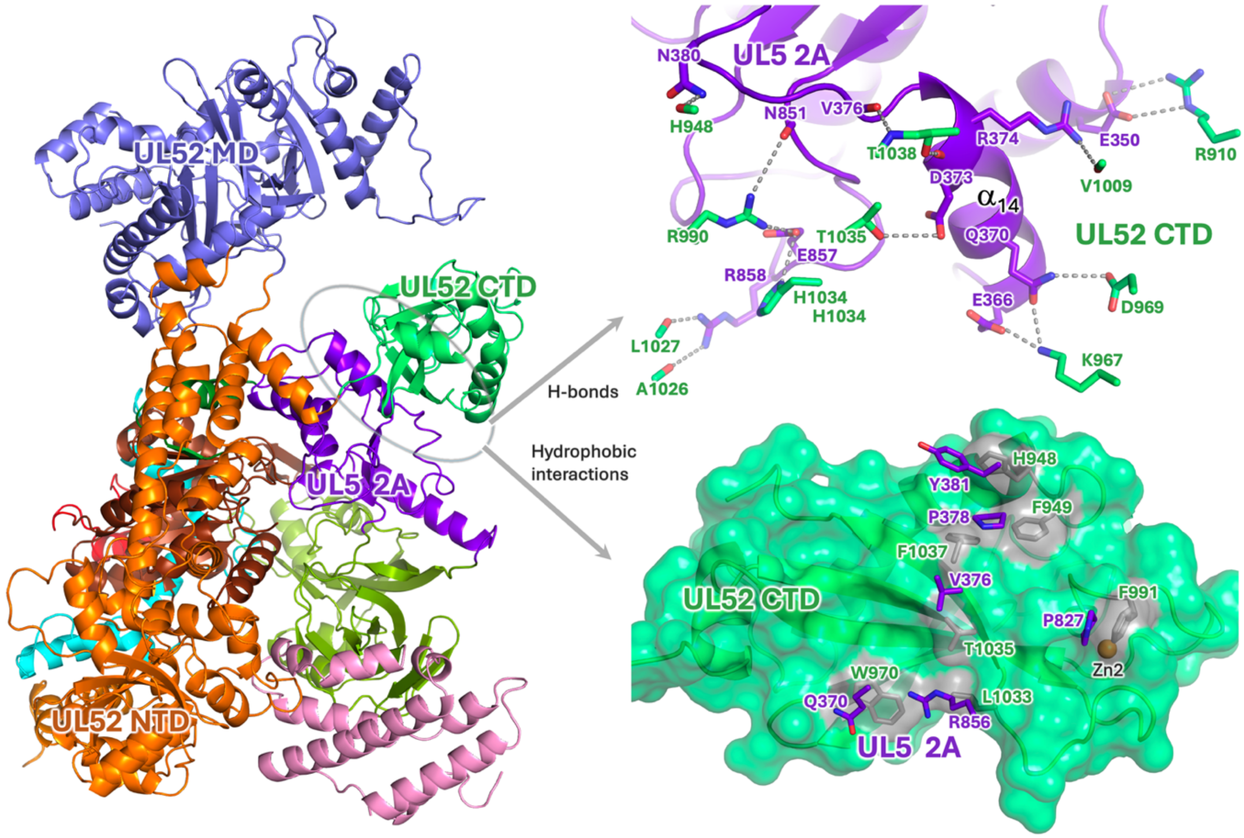

**Figure S17. Interactions between UL5 and CTD of UL52.** The left image shows the location of UL52 CTD relative to UL5 domain 2A. The upper-right image shows inter-subunit H-bonds in detail. The H-bonds are drawn as dashed lines. The lower-right image shows the inter-subunit hydrophobic interactions in detail. The semitransparent surface of UL52 CTD contributing to hydrophobic interactions is highlighted in light grey. In both images on the right, the participating residues are shown as sticks and labeled.

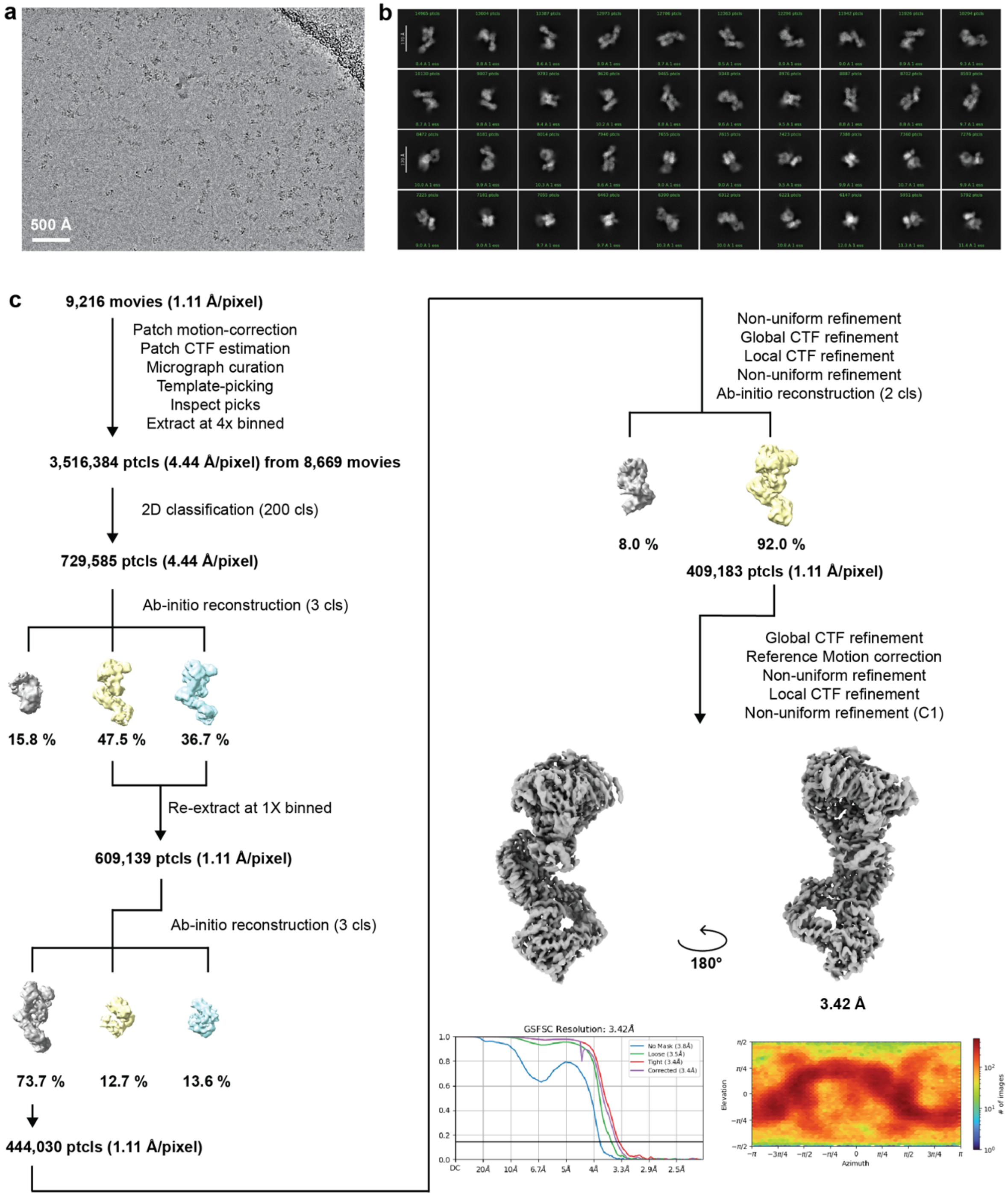

**Figure S18. Cryo-EM processing pipeline for the UL5-UL8-UL52-DNA-pritelivir complex**. (**a**) Representative motion-corrected micrograph (n=9,216) of the cryo-EM dataset for processing. The scale bar dimension is 500 Å. (**b**) 2D classification averages of the UL5-UL8-UL52-DNA-pritelivir complex. A 170 Å scale bar is shown as a white bar. (**c**) The cryo-EM dataset processing pipeline to obtain the consensus map of the UL5-UL8-UL52-DNA-pritelivir complex.

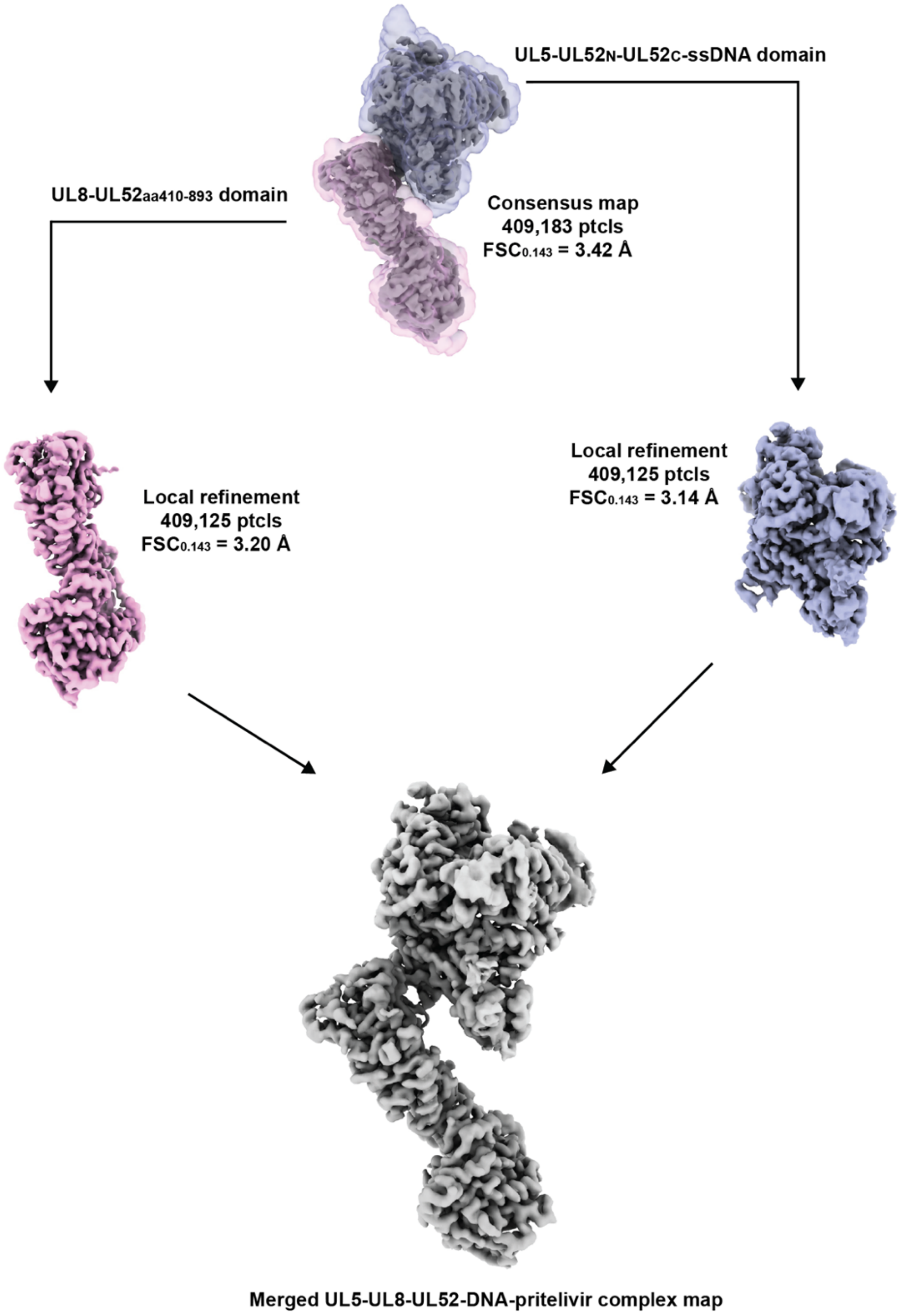

**Figure S19. Local refinement of individual flexible domains of the UL5-UL8-UL52-DNA-pritelivir complex**. The UL5-UL8-UL52-DNA-pritelivir complex was segmented into two distinct domains for particle subtraction and 3D reconstruction. The local masks are shown as semi-transparent. The two domains are merged to yield a better-quality assembled complex map.

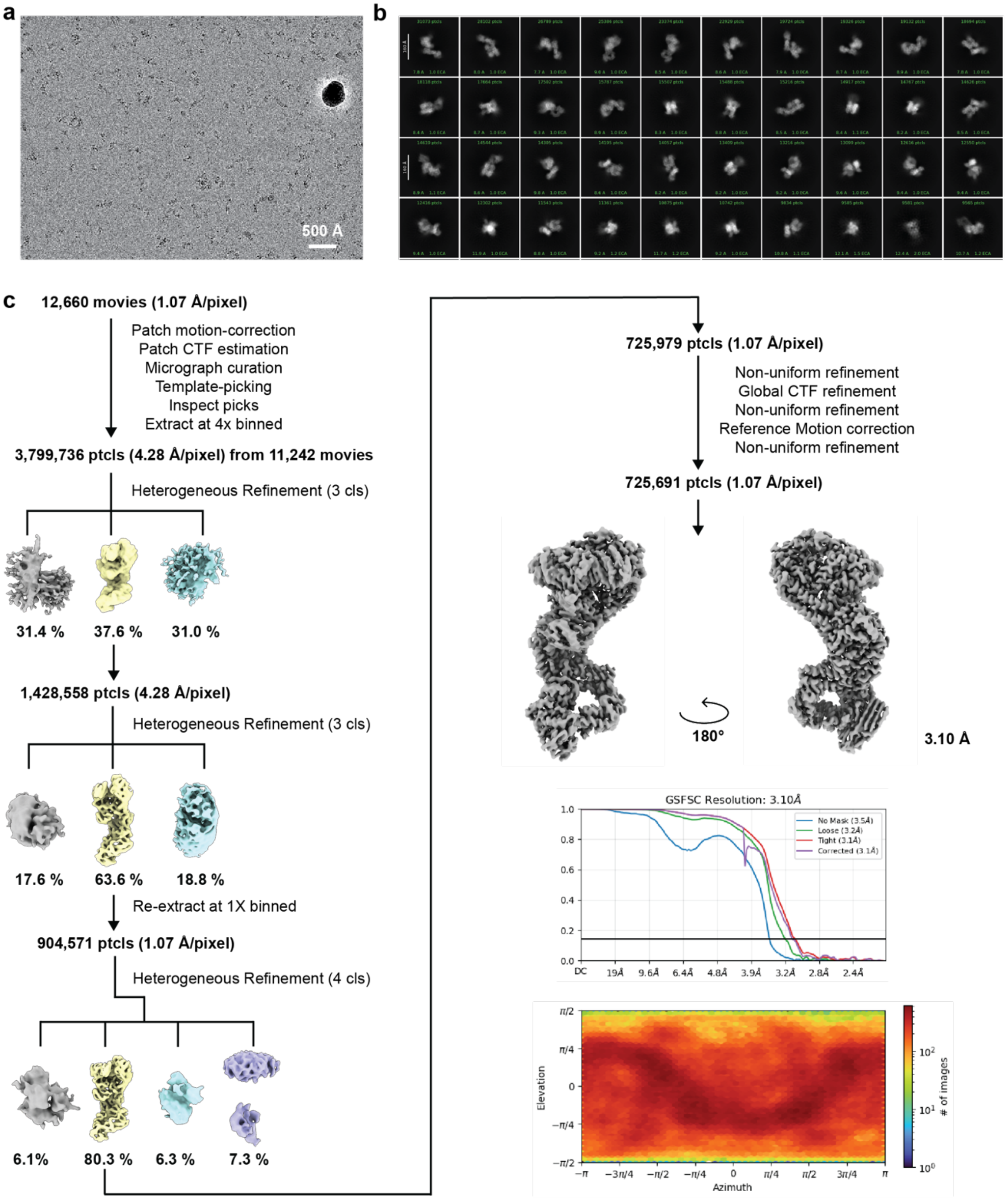

**Figure S20. Cryo-EM processing pipeline for the UL5-UL8-UL52-DNA-amenamevir complex**. (**a**) Representative motion-corrected micrograph (n=12,650) of the cryo-EM dataset for processing. The scale bar dimension is 500 Å. (**b**) 2D classification averages of the UL5-UL8-UL52-DNA-amenamevir complex. A 170 Å scale bar is shown as a white bar. (**c**) The cryo-EM dataset processing pipeline to obtain the consensus map of the UL5-UL8-UL52-DNA-amenamevir complex.

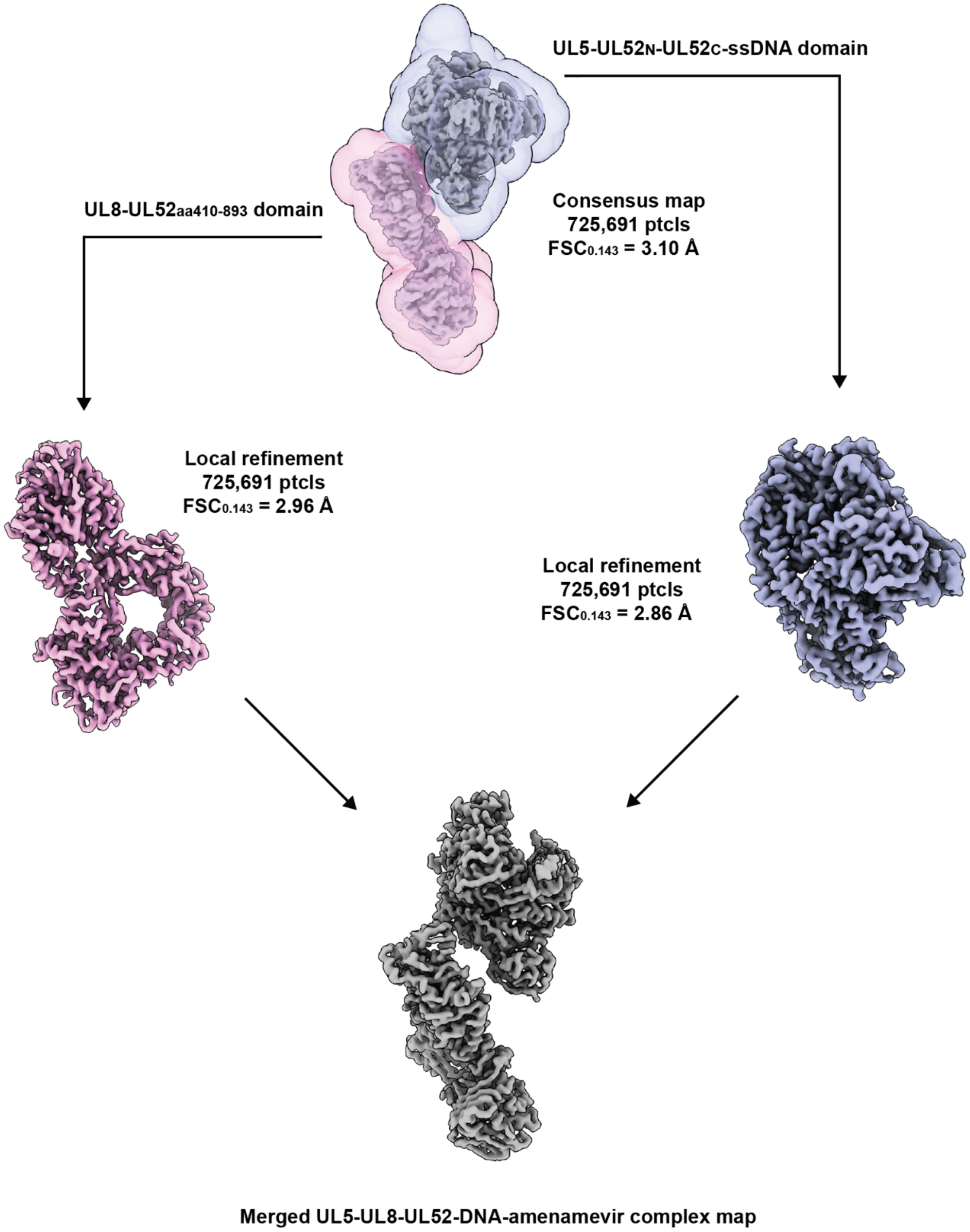

**Figure S21. Local refinement of individual flexible domains of the UL5-UL8-UL52-DNA-amenamevir complex**. The UL5-UL8-UL52-DNA-amenamevir complex was segmented into two distinct domains for particle subtraction and 3D reconstruction. The local masks are shown as semi-transparent. The two domains are merged to yield a better-quality assembled complex map.

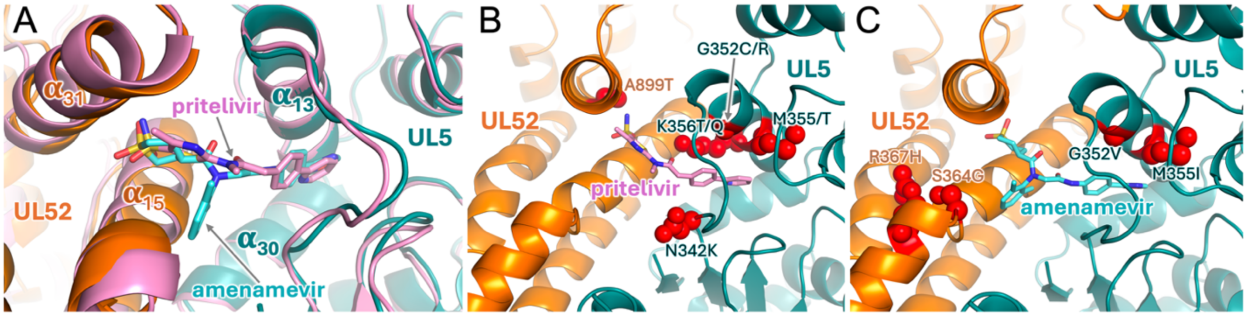

**Figure S22. Comparison of amenamevir and pritelivir binding to HP.** (**A**) Close-up view of the superimposed HP-DNA-amenamevir and HP-DNA-pritelivir complexes. The inhibitors are shown as sticks. (**B**) The structure of HP-DNA-pritelivir. (**C**) The structure of HP-DNA-amenamevir. In the panels (**B**) and (**C**), the side chain atoms of the residues with resistant mutations are displayed as red balls.

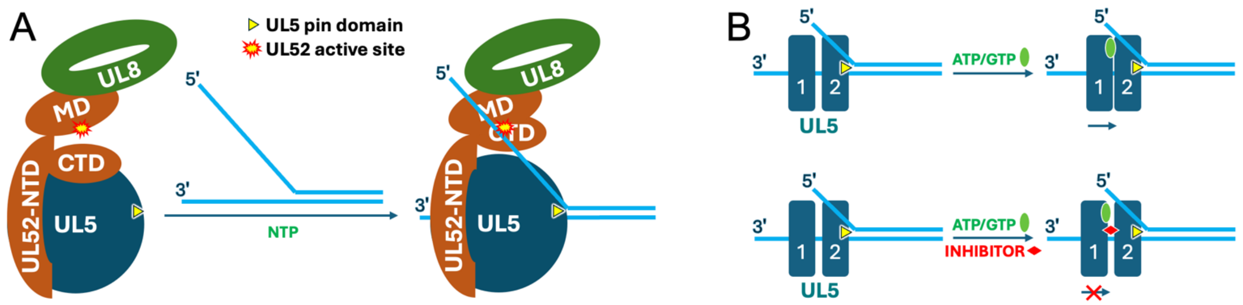

**Figure S23. Cartoon representation of the mechanism of HP function and inhibition.** (**A**) Activation of UL52 primase upon loading on replication fork. The UL52 in apo-HP is not active. Upon loading of HP on replication fork the UL52 CTD is activated by 5'-flap and initiating NTP, resulting in its dissociation from UL5 and then placing the 5'-flap at the primase active site for the initiation of primer's dinucleotide formation. (**B**) Mechanism of UL5 helicase inhibition. In the absence of an inhibitor, ATP/GTP binding results in the movement of UL5 domain 1A in a direction from 3' to 5'. An inhibitor enters between the UL5 domains and, upon ATP/GTP binding, prevents the movement of domain 1A toward domain 2A upon ATP/GTP binding.
